# A fluorescent perilipin 2 knock-in mouse model visualizes lipid droplets in the developing and adult brain

**DOI:** 10.1101/2022.06.21.496932

**Authors:** Sofia Madsen, Ana C. Delgado, Christelle Cadilhac, Fabrice Battison, Vanille Maillard, Elia Magrinelli, Denis Jabaudon, Ludovic Telley, Fiona Doetsch, Marlen Knobloch

**Affiliations:** Department of Biomedical Sciences, University of Lausanne, Lausanne, Switzerland; Biozentrum, University of Basel, Basel, Switzerland; Department of Basic Neurosciences, University of Geneva, Geneva Switzerland; Department of Fundamental Neurosciences, University of Lausanne, Lausanne, Switzerland

## Abstract

Lipid droplets (LDs) are dynamic lipid storage organelles. They are tightly linked to metabolism and can exert protective functions, making them important players in health and disease. Most LD studies *in vivo* rely on staining methods, providing only a snapshot. We therefore developed a LD-reporter mouse by endogenously labelling the LD coat protein perilipin 2 (PLIN2) with tdTomato, enabling staining-free fluorescent LD visualisation in living and fixed tissues and cells. Here we validate this model under standard and high-fat diet conditions and demonstrate that LDs are present in various cells in the healthy brain, including neurons, astrocytes, ependymal cells, neural stem/progenitor cells and microglia. Furthermore, we show that LDs are abundant during brain development and can be visualized using live-imaging of embryonic slices. Taken together, our tdTom-Plin2 mouse serves as a novel tool to study LDs and their dynamics under both physiological and diseased conditions in all tissues expressing Plin2.

## Introduction

Lipids are essential molecules for cells and organisms and serve as energy sources, membrane building blocks and signalling entities. However, the nutritional and cellular availability of lipids fluctuates and too many lipids can be damaging. Thus, organisms have developed specialized intracellular organelles called lipid droplets (LDs) to buffer and regulate the amount of freely available lipids (Walther and Farese, 2012). LDs contain neutral lipids such as triacylglycerols (TAGs) and cholesterol esters and are surrounded by a monolayer of phospholipids and LD coat proteins. While their primary role is to store and regulate access to lipids, it has become clear that besides their metabolic role, LDs are also involved in other important cellular processes such as regulation of cellular stress (Islimye et al., 2022; Jarc and Petan, 2019), protein sequestration (Welte, 2015; Welte and Gould, 2017), inflammation and immune response (Pereira-Dutra and Bozza, 2021).

All cells can form LDs to a certain extent but are under physiological conditions mainly found in lipogenic tissue such as adipose tissue and liver. LDs are tightly linked to lipid metabolism (Walther and Farese, 2012), and can upon nutritional excess be newly formed or increase in size to allow for storage of surplus energy. The liver plays an important role for lipid metabolism, as it both produces and secrets lipids. An imbalance in this process can lead to abnormal accumulation of LDs in the liver, also known as hepatic steatosis (Gluchowski et al., 2017). While liver LD accumulation is a well-known side effect of metabolic diseases, LDs can also accumulate in non-lipogenic tissues e.g. upon obesity or cancer (Cruz et al., 2020; Krahmer et al., 2013). LDs also play an important physiological role in certain non-lipogenic tissues, such as muscle, where they can react dynamically to various stimuli (Seibert et al., 2020). Recent discoveries have put LDs into the spotlight for proper brain functioning (Farmer et al., 2020; Teixeira et al., 2020). Findings in drosophila and mice have linked LD accumulation in the brain to oxidative stress and Alzheimer’s disease (Hamilton et al., 2015; Liu et al., 2015, 2017). Several other studies have shown that lipids can be transferred from neurons to astrocytes, where astrocytes store and subsequently break down lipids stored in LDs. These findings led to the suggestion that there is a metabolic coupling between neurons and astrocytes in order to protect neurons from lipotoxicity (Ioannou et al., 2019; Liu et al., 2017). Moreover, aging and inflammation are important factors shown to trigger LD accumulation in different brain cell types, including neurons, astrocytes, and microglia (Farmer et al., 2020; Marschallinger et al., 2020; Shimabukuro et al., 2016).

LDs have also been described in different brain cells under physiological conditions. They are present in the hypothalamus of both mice and human brains (Maya-Monteiro et al., 2021), and in ependymal cells lining the brain ventricles (Bouab et al., 2011; Capilla-Gonzalez et al., 2014). Furthermore, we recently demonstrated that LDs are important for adult mouse neural stem/progenitor cells (NSPCs) (Ramosaj et al., 2021). NSPCs are prominent during early development and persist throughout adulthood, residing mainly in the subventricular zone (SVZ) and dentate gyrus (DG), where they give rise to new-born neurons and glia (Denoth-Lippuner and Jessberger, 2021; Kriegstein and Alvarez-Buylla, 2009). Both *de novo* lipogenesis (Chorna et al., 2013; Knobloch et al., 2013) and fatty acid beta-oxidation are important pathways for NSPC proliferation and stem cell maintenance and are closely linked to LD build-up and breakdown (Knobloch et al., 2017; Madsen et al., 2021; Stoll et al., 2015; Xie et al., 2017). We have shown that adult mouse NSPCs *in vitro* contain variable amounts of LDs, and that an increase in LDs provides a metabolic context that favours their proliferation (Ramosaj et al., 2021). We also found that LD numbers and sizes changed with NSPC entering quiescence and upon differentiation to neurons and astrocytes (Ramosaj et al., 2021), suggesting that LDs might play a yet unknown role in these processes. Taken together, these findings have led to an increased interest in understanding the role of LDs in brain cells under both physiological and pathological conditions (Farmer et al., 2020; Ralhan et al., 2021; Teixeira et al., 2020).

A current limitation in studying LDs in the brain is their visualisation: Neutral lipid dyes and antibodies against LD coat proteins allow the relatively easy detection of LDs in cells and there are even staining-free microscopy methods to study them *in vitro* (Daemen et al., 2016; DiDonato and Brasaemle, 2003). However, these methods have limitations for studying LD dynamics. We therefore developed an endogenous LD reporter mouse which allows for staining-free fluorescent visualization of LDs in tissues and cells, both living and fixed. To do so, we endogenously tagged the ubiquitously expressed LD marker and coat protein perilipin 2 (Plin2) (Sztalryd and Brasaemle, 2017) with tdTomato using CRISPR/Cas9. We have validated this novel tdTom-Plin2 mouse model under standard and high-fat diet conditions and used it to characterize LDs in the brain.

Using our tdTom-Plin2 mouse, we show that LDs are abundant in the brain from embryonic development to adulthood and can be found in the main brain cell types such as neurons, astrocytes, microglia and oligodendrocytes as well as ependymal cells, endothelial cells, NSPCs and their progeny. We furthermore show that cells can be isolated from these mice and sorted based on LD content using tdTomato and used for subsequent analysis such as single cell RNA sequencing (scRNA seq). Importantly, we also demonstrate that tissues from these mice can be used for live imaging, enabling to study LD dynamics in complex set-ups. As *Plin2* is ubiquitously expressed, the tdTom-Plin2 mouse will serve as a powerful novel tool to study LDs in different types of tissues during both physiological and pathological conditions.

## Results

### Generation of the endogenous LD reporter mouse and validation of tdTomato-tagged PLIN2 in NSPCs

To visualize LDs in a staining-free manner, we generated a novel LD reporter mouse by endogenously tagging the LD-specific and ubiquitously expressed PLIN2 protein with tdTomato. We inserted *TdTomato* with a short linker sequence at the N-terminus of *Plin2* gene in mouse embryonic stem cells (ESCs) using CRISPR/Cas9 (Fig. 1A and B). Correctly edited ESCs were injected into mouse blastocysts to generate chimeras, and a germline transmitting tdTom-Plin2 mouse line was established (Fig. 1B).

**Figure 1.**
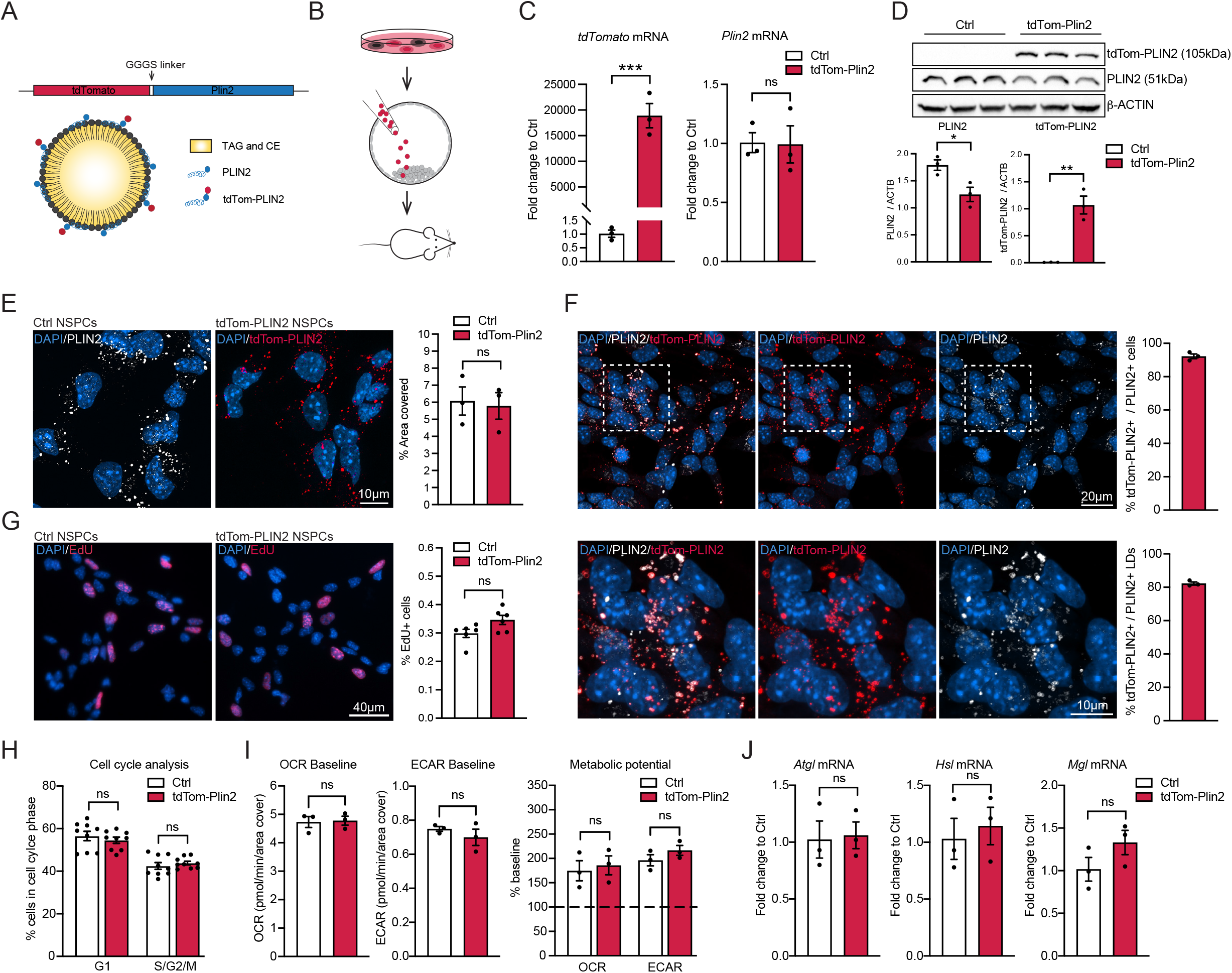
Generation of the endogenous LD reporter mouse and validation of tdTomato-tagged PLIN2 in NSPCs. **A)** Schematic illustration of the construct used to generate the endogenous LD reporter mouse. TdTomato is linked to the N-terminus of *Plin2* on one allele, which results in LDs coated with wt PLIN2 and tdTomato-tagged PLIN2. **B)** Generation of the tdTom-Plin2 mouse. Mouse ESCs were CRISPRed *in vitro*. Correctly edited ESC clones were injected to blastocysts to generate chimeras, which were then screened for germ-line expression. **C)** Analysis of mRNA expression by RT-qPCR show *tdTomato* expression in NSPCs from the tdTom-Plin2 mouse and no significant difference in *Plin2* between wt NSPCs and tdTom-Plin2 NSPCs. (n=3 samples per condition, fold change +/- SEM). **D)** Western blot analysis shows the presence of PLIN2 and tdTom-PLIN2 in NSPCs from the tdTom-Plin2 mice, whereas only untagged PLIN2 is found in wt NSPCs. **E)** The area covered by PLIN2 is comparable in wt NSPCs (stained against PLIN2) and in tdTom-Plin2 NSPCs (revealed by tdTom-PLIN2). (n=3 coverslips per condition, 237 and 233 cells respectively, mean +/- SEM). **F)** Quantification shows that > 95 % of the NSPCs from the tdTom-Plin2 mouse express tdTom-PLIN2 (n= 3 coverslips per condition, 233 cells, +/- SEM) and 80 % of the PLIN2+ LDs are tdTom-PLIN2+ in NSPCs from the tdTom-Plin2 mouse. (n= 3 coverslips per condition, 222 cells, +/- SEM). **G)** Quantification of EdU positive cells shows no significant difference in proliferation between wt and tdTom-Plin2 NSPCs. (n=6 coverslips per condition, mean +/- SEM). **H)** Cell cycle analysis by flow cytometry confirms that there is no difference in NSPC proliferation due to the tagged PLIN2. (n= 9 samples per condition, mean +/- SEM). **I)** Basic metabolic properties are similar between Ctrl and tdTom-Plin2 NSPCs. Bar graphs show baseline oxygen consumption rate (OCR) and extracellular acidification (ECAR), as well as the metabolic potential (n=3 experiments, 12-15 replicates per experiment and condition, mean +/- SEM) **J)** The three major LD lipases, adipose triglyceride lipase (*Atgl*), hormone sensitive lipase (*Hsl*) and monoacylglycerol lipase (*Mgl*) are similarly expressed in wt and tdTom-Plin2 NSPCs as shown by RT-qPCR (n=3 samples per condition, fold change +/- SEM). Asterisks indicate the following p-values: * < 0.05; ** < 0.01; *** < 0.001. ns= non-significant.

We have recently shown that primary NSPCs from adult mice have numerous PLIN2-positive LDs *in vitro* (Ramosaj et al., 2021). We therefore used primary NSPCs, isolated from heterozygous tdTom-Plin2 mice, to validate this novel LD reporter model *in vitro*. We could detect *TdTomato* mRNA in tdTom-Plin2 NSPCs, and they expressed similar levels of *Plin2* mRNA as control (Ctrl) NSPCs (Fig. 1C). Western blot analysis showed a shift in PLIN2 size due to the heterozygous tdTomato tag in NSPCs from tdTom-Plin2 mice and a reduction in untagged PLIN2 levels compared to Ctrl NSPCs (Fig. 1D). Quantification of the endogenous tdTom-PLIN2 signal in tdTom-Plin2 NSPCs showed a similar area coverage as staining against PLIN2 in Ctrl NSPCs (Fig.1E), and 95 % of the NSPCs expressing PLIN2 also expressed tdTom-PLIN2 and 80% of all LDs expressing PLIN2 showed good colocalization with the endogenous tdTom-PLIN2 (Fig. 1F).

Having previously shown that LDs influence NSPC proliferation and metabolism (Ramosaj et al., 2021), we next assessed if the tdTom-Plin2 tag might alter these two characteristics. EdU pulse-labelling of proliferating cells showed no significant changes in NSPC proliferation in tdTom-Plin2 versus Ctrl NSPCs (Fig. 1G). Using flow cytometry-based cell cycle analysis, we confirmed these results, showing that the tag did not alter NSPC proliferation (Fig. 1H). Ctrl and tdTom-Plin2 NSPCs had similar baseline oxygen consumption (OCR) and extracellular acidification (ECAR), as well as a similar metabolic potential, demonstrating that the tag did not alter basic metabolic properties (Fig. 1I). As tagging a LD coat protein bears the risk of influencing LD accessibility, we also assessed the mRNA expression of the three major LD lipases, Adipose triglyceride lipase (*Atgl*), Hormone sensitive lipase (*Hsl*) and Monoacylglycerol lipase (*Mgl*). We found no statistically significant differences when comparing NSPCs from tdTom-PLIN2 and Ctrl mice (Fig. 1J). Taken together, these data show that the endogenous tdTom-PLIN2 signal labels LDs and that the tdTomato tag does not affect the overall expression of *Plin2*, LD abundance, proliferation, nor basic metabolic properties in NSPCs.

### TdTom-Plin2 mice show normal weight gain, normal fat accumulation and report LDs in various organs

Overexpression of PLIN2 has been reported to attenuate lipolysis by reducing the access of adipose triglyceride lipase (ATGL), thus stabilizing LDs (Listenberger et al., 2007). Having endogenously tagged *Plin2*, our reporter system does not rely on overexpression. However, to rule out that the endogenous tag on one allele might affect LD stability and fat accumulation, we monitored the weight progression from the age of 3 weeks to 8 weeks of tdTom-Plin2 heterozygous and Ctrl littermates *in vivo* and found no major differences in weight gain (Fig. 2A). A more detailed fat composition analysis using EchoMRI showed no significant differences in fat mass when comparing 8-week-old tdTom-Plin2 and Ctrl mice (Fig. 2B), thus the tdTomato tag does not seem to influence the body weight and body fat composition of this novel reporter mouse.

**Figure 2.**
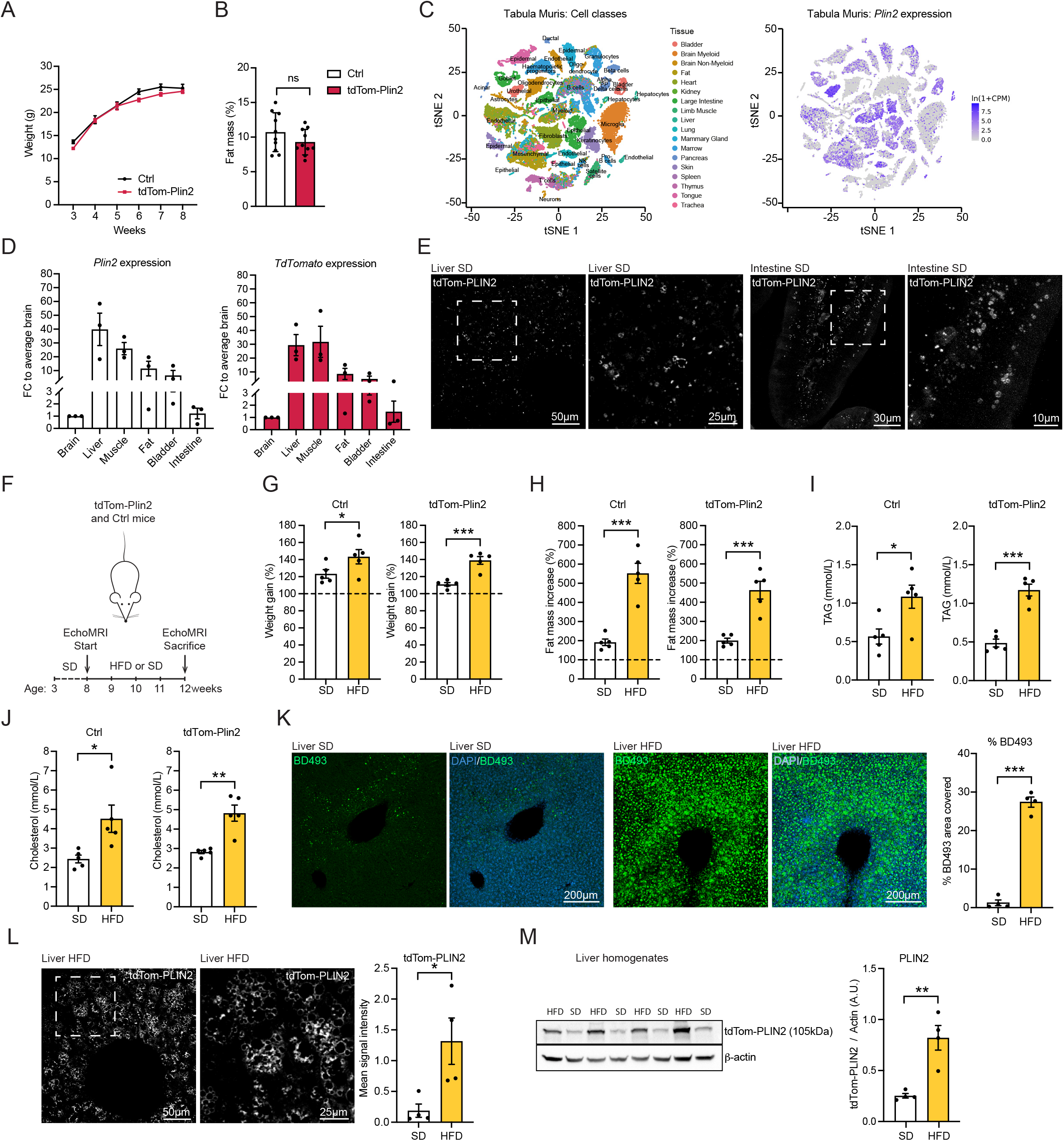
TdTom-Plin2 mice report LDs in various organs, increase fat mass and increase LDs in the liver upon a short-term high fat diet. **A)** There is no major difference in body weight gain over time between Ctrl and tdTom-Plin2 mice between 3- and 8 weeks of age. (n=10 mice per group). **B)** Assessment of body fat composition at 8 weeks using EchoMRI shows no significant difference between Ctrl and tdTom-Plin2 mice. (n=10 mice per group, mean +/- SEM). **C)** *Plin2* is expressed in many tissues such as liver, lung, and heart, as shown by a query of the *Tabula muris* (Schaum et al., 2018). **D)** *Plin2* and *tdTomato* are expressed in many organs in the tdTom-Plin2 mouse (n=3 mice, mean +/- SEM). **E)** Representative images of the endogenous tdTomato signal from the liver and the intestine of a tdTom-Plin2 mouse confirm the presence of fluorescent LDs in these organs. **F)** Overview of the HFD exposure. **G)** Both tdTom-Plin2 and Ctrl mice had a significant increase in body weight after 4 weeks of HFD compared to SD. (n=5 mice per group, mean +/- SEM). **H)** Both tdTom-Plin2 and Ctrl mice had a significant increase in fat mass after 4 weeks of HFD compared to SD. (n=5 mice per group, mean +/- SEM). **I** and **J)** Serum levels of TAGs and cholesterol were significantly increased in HFD fed animals compared to SD in both tdTom-Plin2 and Ctrl mice. (n=5 mice per group, mean +/- SEM). **K)** Liver LDs, revealed by BODIPY 493/503 (BD493), accumulated significantly in tdTom-Plin2 mice exposed to a HFD. **L)** Similarly, endogenous tdTom-PLIN2 increased as well significantly in tdTom-Plin2 mice fed a HFD. **M)** TdTom-PLIN2 protein was significantly increased in HFD liver homogenates, as revealed by western blot analysis. Asterisks indicate the following p-values: * < 0.05; ** < 0.01; *** < 0.001. ns= non-significant.

*Plin2* is expressed in many tissues such as liver, lung, and heart (Fig. 2C), as shown by a query of the *Tabula muris*, a single cell RNA sequencing (scRNA seq) database of cells from 20 different mouse organs (Schaum et al., 2018). RT-qPCR confirmed that tdTomato-tagged *Plin2* was expressed in various organs of heterozygous tdTom-Plin2 mice (Fig. 2D). Imaging endogenous tdTom-PLIN2 by confocal microscopy furthermore showed clear ring-like signal in tissue sections of liver and intestine from tdTom-Plin2 mice (Fig. 2E). These data show that the endogenously tagged Plin2 is expressed in several organs, thus making the tdTom-Plin2 mouse a usable tool for studying LDs in different types of tissues.

### Exposure to a short-term high fat diet increased fat mass and accumulation of tdTom-Plin2 in the liver of TdTom-Plin2 mice

We next wanted to assess this novel endogenous LD reporter mouse under altered lipid availability. We therefore exposed tdTom-Plin2 mice and Ctrl mice to a short-term high fat diet (HFD), to increase the circulating lipids but limit the manifold side-effects of prolonged HFD such as severe obesity and diabetes. Adult heterozygous tdTom-Plin2 and Ctrl mice were either fed with a HFD containing 60% fat or a standard chow diet (SD) for 4 weeks. We monitored the body weight weekly and performed EchoMRI measurements at the beginning and end of the experiment (Fig. 2F). Both Ctrl and tdTom-Plin2 mice on HFD had a 40 % weight increase over 4 weeks, whereas the SD fed control groups only had a 10-20 % weight gain (Fig. 2G). Similarly, the fat mass increased more than 4 times in both Ctrl and tdTom-Plin2 mice on HFD, whereas SD fed mice only had a 2-fold increase in fat mass over 4 weeks (Fig. 2H). Moreover, we collected serum at the end of the experiment to assess how the mice reacted to the diet. Mice fed with HFD had significantly higher serum levels of TAGs and cholesterol (Fig. 2I and 2J, S1A and S1B), confirming that HFD indeed increased the levels of circulating lipids. We did not observe any differences in free fatty acids (Fig. S1C). Blood levels of alanine aminotransferase (ALAT) and aspartate aminotransferase (ASAT), which are used as indicators of liver disease, were not significantly changed with HFD, even though tdTom-Plin2 mice on HFD had a slight elevation in both (Fig. S1D and S1E). Taken together, these data show that tdTom-Plin2 mice react in a similar way to increased lipid availability as Ctrl mice, with increased fat mass and weight gain. Thus, endogenously tagging *Plin2* does not have an overall effect on lipid storage in the LD reporter mice.

We next examined the livers of tdTom-Plin2 mice, as it has previously been shown that the liver reacts to increased circulating lipids with increased LD number and size (Gluchowski et al., 2017). Indeed, quantification of the total area covered by the neutral lipid dye BODIPY493/503 (BD493) showed a significant increase when exposed to a HFD (Fig. 2K). Similarly, endogenous tdTom-PLIN2 also increased significantly in tdTom-Plin2 mice fed with a HFD (Fig. 2L, S1F). Confirming these data, tdTom-PLIN2 protein was significantly increased in HFD liver homogenates, as revealed by western blot analysis (Fig. 2M). These data show that lipid availability alters LDs in the liver and that the endogenous tdTomato signal reports such changes.

### LDs are present in multiple cell types in the healthy adult mouse brain

The presence of LDs in the healthy mammalian brain has to date only been described in few cell types. As the tdTom-Plin2 mouse reports LDs in isolated NSPCs (Fig. 1) and in the liver and intestine (Fig. 2), we next explored if the tdTom-Plin2 mouse would be a good tool to visualize and characterize LDs in the postnatal and adult brain. We first measured the expression of *Plin2* in whole brain extracts of 2-month-old tdTom-Plin2 and Ctrl mice. In line with the data obtained with NSPCs *in vitro*, *tdTomato* mRNA expression was detectable in tdTom-Plin2 mice, and *Plin2* mRNA was expressed in a similar manner as in Ctrl brains (Fig. 3A). Expression of the main lipases *Atgl*, *Hsl* and *Mgl* was also comparable between Ctrl and tdTom-Plin2 mice (Fig. S2A), although *Atgl* seemed slightly less expressed in tdTom-Plin2 brains. Moreover, lipidomic analysis showed that the levels of TAGs, diacyglycerides (DAGs) and monoacylglycerols (MAGs) were similar between brains from tdTom-Plin2 and Ctrl mice (Fig. S2B). This suggests that the tdTomato tag does not significantly alter neutral lipids in the brain. Confocal microscopy of unstained sagittal brain sections from 2-month-old tdTom-Plin2 mice revealed a surprising abundance and size diversity of fluorescently labelled endogenous LDs in various brain regions, such as the olfactory bulb, cortex, the lateral wall of the lateral ventricles and cerebellum (Fig. 3B and 3C). To validate that this tdTomato signal indeed reports LDs *in vivo*, we focused on the lateral wall, where studies have reported large LDs in ependymal cells (Ralhan et al., 2021). TdTomato positive rings of various sizes were clearly present in the lateral wall confirming that these were LDs by co-staining against PLIN2 (Fig. 3D).

**Figure 3.**
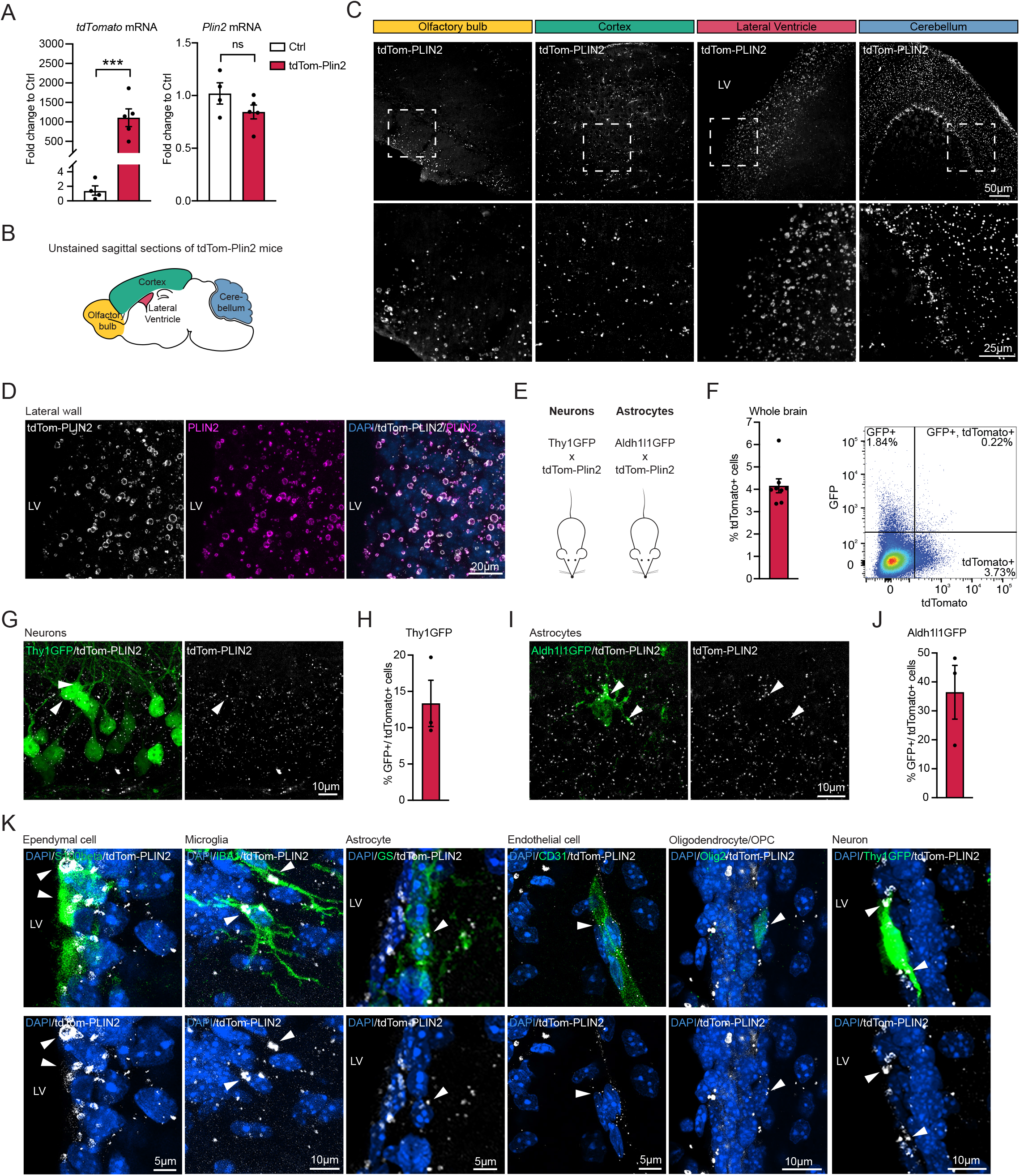
LDs are present in multiple cell types in the healthy adult mouse brain. **A)** *TdTomato* expression is clearly detectable in tdTom-Plin2 brains, and *Plin2* mRNA is expressed in a similar manner as in Ctrl brains. (n=4-5 mice per group, fold change +/- SEM). **B)** Schematic illustration of a sagittal brain section, highlighting specific regions **C)** Endogenous tdTom-PLIN2 signal in the olfactory bulb, cortex, lateral ventricle well and cerebellum shows abundant LDs in the brain. Representative maximum intensity projections at low (top) and high (bottom) magnifications. **D)** TdTomato positive rings of various sizes are clearly present in the lateral ventricle wall and colocalize with the signal from staining against PLIN2, confirming that they are LDs. Representative maximum intensity projection of individual channels and a merge. **E)** Scheme of the crossing of tdTom-Plin2 mice to Thy1GFP labelling neurons, and Aldh1l1GFP reporter mice, labelling astrocytes. **F)** FACS analysis of whole brain single cell suspensions shows that about 4 % of the cells in the mouse brain express tdTom-Plin2, thus have LDs. (n=8 mice, +/- SEM). Representative FACS plot. **G)** FACS analysis of whole brain cell extraction from the tdTom-Plin2 mice crossed to Thy1GFP mice show 13 % of all Thy1GFP positive neurons have LDs. (n=3 mice, mean +/- SEM). **H)** FACS analysis of whole brain cell extraction from the tdTom-Plin2 mice crossed to Aldh1l1GFP mice show that around 40 % of all Aldh1l1GFP positive astrocytes have LDs. (n=3 mice, mean +/- SEM). **I)** Immunohistochemical staining of the lateral wall show a large variability of endogenous tdTom-PLIN2 positive LDs in S100b-positive ependymal cells, IBA1-positive microglia, GS-positive astrocytes, CD31 positive endothelial cells, OLIG2-positive oligodendrocytes/oligodendrocyte precursors and a Thy1GFP neuron. Representative maximum intensity projections of observed cell types. Asterisks indicate the following p-values: *** < 0.001. ns= non-significant.

To elucidate which cell types that contained tdTom-PLIN2 positive LDs under physiological conditions, we crossed tdTom-Plin2 mice to two different GFP-reporter lines (Fig. 3E), labelling neurons (Thy1GFP, (Feng et al., 2000)) and astrocytes (Aldh1l1-GFP, (Gong et al., 2003)). For a global quantitative view, we first isolated cells from whole brains and analysed them by FACS, revealing that around 4% of all viable brain cells express tdTomato, thus have LDs (Fig. 3F and S2C). Cell-type specific analyses showed that around 13% of the Thy1GFP positive neurons (Fig. 3G and S2C) and around 40% of the Aldh1l1GFP positive astrocytes had tdTomato positive LDs (Fig. 3H and S2D).

To further characterise cell types that contain LDs, we used immunohistochemistry to stain for different cell type markers. Imaging of the lateral wall showed endogenous tdTomato expression in ependymal cells (S100 Calcium Binding Protein B, S100B), microglia (ionized calcium binding adaptor molecule 1, IBA1), astrocytes (glutamine synthetase, GS), endothelial cells (cluster of differentiation 31, CD31), and oligodendrocytes/oligodendrocyte precursors (oligodendrocyte transcription factor 2, OLIG2) (Fig. 3I). We also detected LDs in Thy1 positive neurons in the lateral wall of the ventricle in the Thy1GFP tdTom-Plin2 reporter mice (Fig. 3I). Taken together, these data show that LDs are present in multiple cell types in the mouse brain under physiological conditions and to a larger extent that previously thought.

### ScRNA sequencing of tdTomato positive microglia reveals a subpopulation with a specific signature under physiological conditions

One important advantage of the tdTom-Plin2 mouse is the staining-free detection of LD-containing cells due to the endogenous fluorescence. This allows for separation of living cell populations based on whether they contain LDs or not and for subsequent characterisation of signature features of such populations. LD-accumulating microglia (LDAM) with a proinflammatory gene expression signature have been identified in the aging brain of mice and humans using BD493 staining (Marschallinger et al., 2020). Given that we already detect tdTom-Plin2 positive LDs in microglia in 2-month old tdTom-Plin2 mice (Fig. 3I), we explored if we could detect a similar signature distinguishing tdTomato positive and tdTomato negative microglia. We used whole brain cell extractions and FACS to separate cells based on tdTomato expression, then performed scRNA seq on the tdTomato positive and negative fractions, focusing on microglial clusters (Fig. 4A). Cells with microglial identity clustered into three distinct clusters (Fig. 4B and Fig. S3A). Interestingly, the three clusters contained distinct proportions of tdTomato positive cells (Fig. 4C). Cluster 0 contained comparable numbers of tdTomato positive and tdTomato negative cells (56% vs 44%), cluster 1 had lower proportion of tdTomato positive cells (24% vs 76%) whilst cluster 2 was almost entirely composed of tdTomato positive cells (95% to 5%) (Fig. 4D). The *Plin2* expression matched this distribution, validating the sorting approach (Fig. 4E). A heatmap representation based on the top differentially expressed genes per cluster showed a clear separation of the tdTomato positive cluster 2, whereas cluster 1 and 0 were more similar (Fig. 4F). To better understand the cluster 2 specific signature, we compared differentially expressed genes between cluster 1 (mainly tdTomato negative cells) and cluster 2 (mainly tdTomato positive cells). 395 genes were significantly upregulated in the tdTomato positive cluster 2, and 153 significantly downregulated (Fig. 4G). A gene ontology (GO) analysis of the upregulated genes revealed the enrichment of several terms involved in cellular activation (e.g. GO: 0006412: translation; GO:0050865: regulation of cell activation) as well as in immune functions (e.g. GO:0001817: regulation of cytokine production; GO:0002252: immune effector process; GO:0001774: microglial cell activation). Together, this analysis suggests that tdTomato positive microglia might represent a fraction of microglia undergoing activation. Previous studies have identified microglial signatures related to disease states (disease-associated microglia, DAM) and a recent review summarized the most common microglial gene expression alterations across various neurodegenerative diseases (Chen and Colonna, 2021). We queried whether these genes differed between our clusters 1 and 2. Indeed, many genes upregulated with disease were also expressed more highly in tdTomato positive microglia (Fig. S3B). However, so-called homeostatic microglial genes which are usually downregulated with disease, were also expressed more highly in the tdTomato positive microglia (Fig. S3B), suggesting that these microglia might be prone to adapt a DAM-like signature, but are not yet in the same state. Similarly, the comparison with the most significantly up-and downregulated LDAM genes (Marschallinger et al., 2020) showed that tdTomato positive microglia share some of the LDAM features but are not entirely similar (Fig. S3C). Whether microglia will shift their gene expression profile with aging or disease remains to be determined.

**Figure 4.**
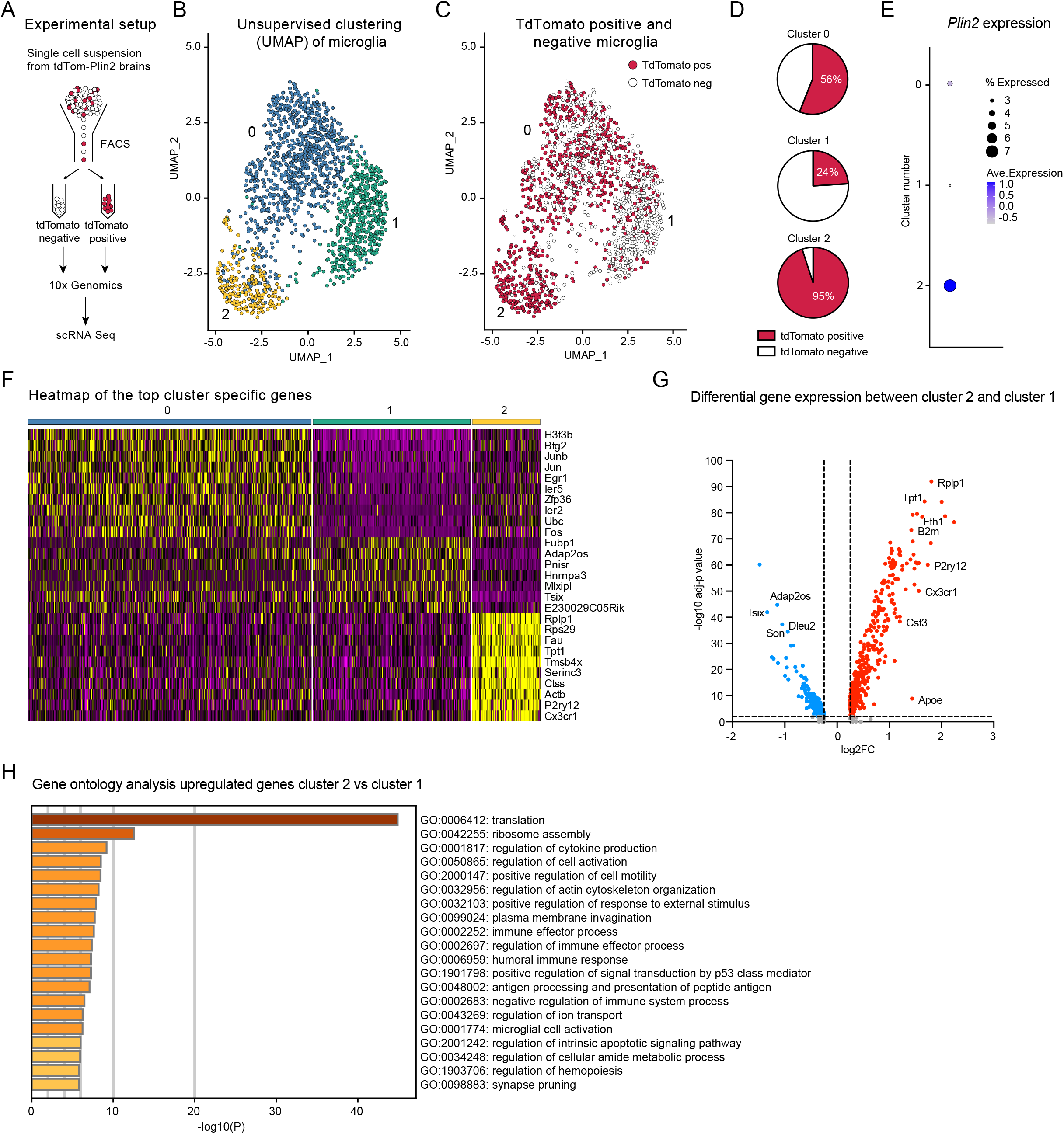
ScRNA sequencing of tdTomato positive microglia reveals a subpopulation with a specific signature under physiological conditions. **A)** Schematic illustration of the experimental setup. **B)** Unsupervised clustering of cells with microglial identity reveals three distinct clusters. **C)** The three microglial clusters contain distinct proportions of tdTomato positive cells. **D)** Quantification of tdTomato positive and negative cells per cluster identifies cluster 2 as highly enriched for microglia containing LDs, while cluster 1 contains a majority of tdTomato negative cells. **E)** The *Plin2* expression matches this distribution, validating the sorting approach. **F)** A heatmap of the top cluster specific genes shows a clear separation of the tdTomato positive cluster 2, whereas cluster 1 and 0 seem to be more similar. **G)** Volcano plot of differentially expressed genes between cluster 2 and cluster 1. **H)** A gene ontology (GO) analysis of the upregulated genes in cluster 2 reveals the enrichment of several terms involved in cellular activation as well as in immune functions.

Taken together, we demonstrate that the tdTom-Plin2 mouse can be used to identify and characterize subpopulations of cells based on their LD content. Importantly, we show here a proof-of-concept with microglia, but this should be possible for any cell type than can be extracted and subsequently sorted by FACS.

### LDs are present in NSPCs and their progeny in the postnatal and adult mouse brain

We next sought to characterise LDs in NSPCs *in vivo* using tdTom-Plin2 mice crossed with Nestin-GFP mice (Nes-GPF, (Yamaguchi et al., 2000)). Nes-GFP labels the NSPCs in the dentate gyrus (DG) and the NSPCs and ependymal cells in the subventricular zone (SVZ) (Fig. 5A). TdTomato positive LDs were detected in both the DG and the SVZ in brains of 1-, 3- and 8-week-old mice (Fig. 5B and C, S4A and S4B). We performed FACS analysis of micro-dissected SVZ and hippocampus of 8-week-old NesGFP-mice to assess the numbers of double positive cells. Around 30% of the NesGFP positive NSPCs in the hippocampus were tdTomato positive (Fig. 5D and S4C) respectively 13% of the NesGFP positive cells in the SVZ were tdTomato positive (Fig. 5E and S4D). However, since NesGFP is also partially expressed in ependymal cells (Yamaguchi et al., 2000), we cannot define the exact percentage of NSPCs in the SVZ that have LDs. In summary, we show that NSPCs *in vivo* have LDs, but not to the same extent as *in vitro* where almost all NSPCs have LDs (Fig. 1E) (Ramosaj et al., 2021).

**Figure 5.**
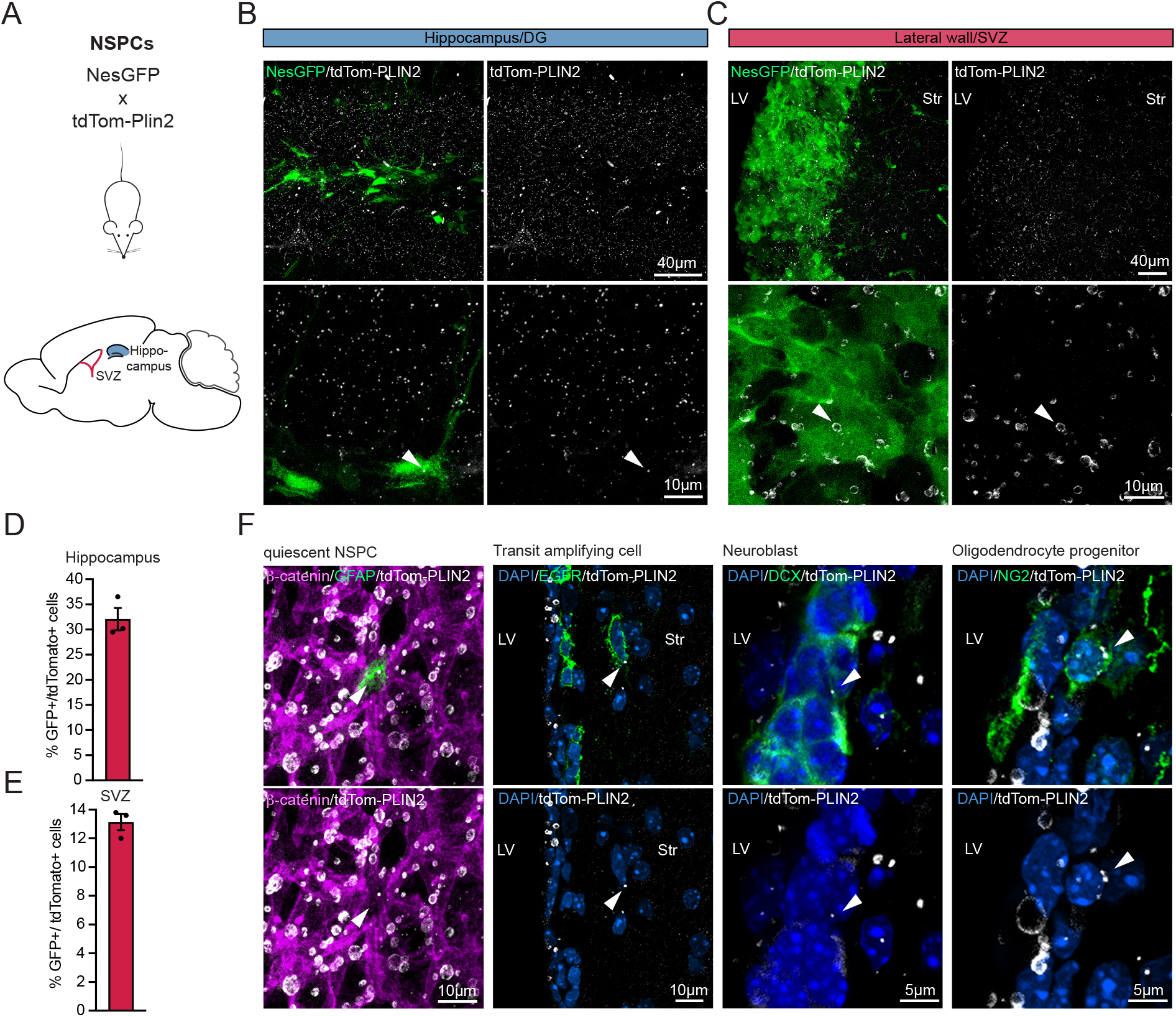
LDs are present in NSPCs and their progeny in the postnatal and adult mouse brain. **A)** Scheme of the crossing of tdTom-Plin2 mice to NesGFP reporter mice, labelling NSPCs in the SVZ and DG. **B** and **C)** Endogenous tdTom-PLIN2 signal in the DG and SVZ showing LDs in NSPCs. Representative images of non-stained sections show NesGFP positive NSPCs and tdTom-PLIN2 positive LDs. (Maximum intensity projections,10 μm stacks).**D)** FACS analysis of cells isolated from the hippocampus of tdTom-Plin2 mice crossed to NesGFP mice show that around 30 % of the NesGFP positive NSPCs have LDs. (n=3 mice, mean +/- SEM). **E)** FACS analysis of cells isolated from the SVZ of tdTom-Plin2 mice crossed to NesGFP mice show that around 13 % of the NesGFP positive NSPCs have LDs. (n=3 mice, mean +/- SEM). **F)** Immunohistochemical staining of cells in the SVZ show endogenous tdTom-PLIN2 positive LDs in GFAP positive quiescent NSPCs, EGFR positive transit amplifying cells, DCX positive neuroblasts and NG2 positive oligodendrocyte progenitors. Representative maximum intensity projections of the observed cell types.

To further characterize LDs in NSPCs and their progeny, we immunostained for different markers of the NSPC lineage in the SVZ. We detect LDs in some, but not all, glial fibrillary acidic protein (GFAP) expressing radial stem cells, epidermal growth factor receptor (EGFR) expressing transit amplifying cells, doublecortin (DCX) expressing neuroblast and neuron-glial antigen 2 (NG2) expressing oligodendrocyte progenitors (Fig. 5F). Taken together, our endogenous LD reporter mouse reveals that LDs are present in both NSPCs and their progeny in the adult mouse brain under physiological conditions.

### Short-term high fat diet increases the total TAG levels in the brain and leads to slightly increased LDs in the wall of the lateral ventricles

We have previously shown that supplement of exogenous lipids *in vitr*o increases LDs in NSPCs (Ramosaj et al., 2021). Additionally, Hamilton and colleagues showed that infusion of oleic acid (OA) directly into the lateral ventricle of adult mice using an osmotic pump led to an increase of LDs in the ependymal layer of the SVZ (Hamilton et al., 2015). We therefore investigated if a short term HFD affects LDs in the mouse brain. We first assessed gene expression of *Plin2* and *tdTomato* in brains from tdTom-Plin2 mice and Ctrl mice which had been fed with a SD or short term HFD (Fig. 2E).

Neither *Plin2* nor *tdTomato* were significantly changed due to the diet (Fig. 6A and B), indicating that *Plin2* expression does not increase in the brain upon short term HFD. However, changes in LDs could be local, and therefore difficult to detect at the global mRNA level from the whole brain. As LDs mainly store TAGs, we performed quantitative lipidomics on the same brains, a highly sensitive method providing a detailed analysis of the neutral lipid composition in the brain. The lipidomics revealed that HFD fed mice had an increase in TAGs, reaching statistical significance for the Ctrl mice and a trend in the tdTom-Plin2 mice (Fig. 6C). When looking at the TAG levels independently from genotype, HFD significantly increased TAG levels in the brain (Fig. 6D). The precursors of TAGs, DAGs and MAGs, were not significantly changed in any of the groups (Fig. S5A and S5B). A more detailed analysis of the lipid composition in the brain of HFD compared to SD fed mice showed that 18% of the measured lipids were significantly changed in either Ctrl or tdTom-Plin2 mice, or in both (Fig. 6E). The majority of the significantly changed brain lipids were TAGs (Fig. 6F). When looking at the fold changes of the significantly changed lipids, both Ctrl and tdTom-Plin2 brains showed a coherent pattern in terms of increased and decreased lipid species as a reaction to the HFD (Fig. 6F).

**Figure 6.**
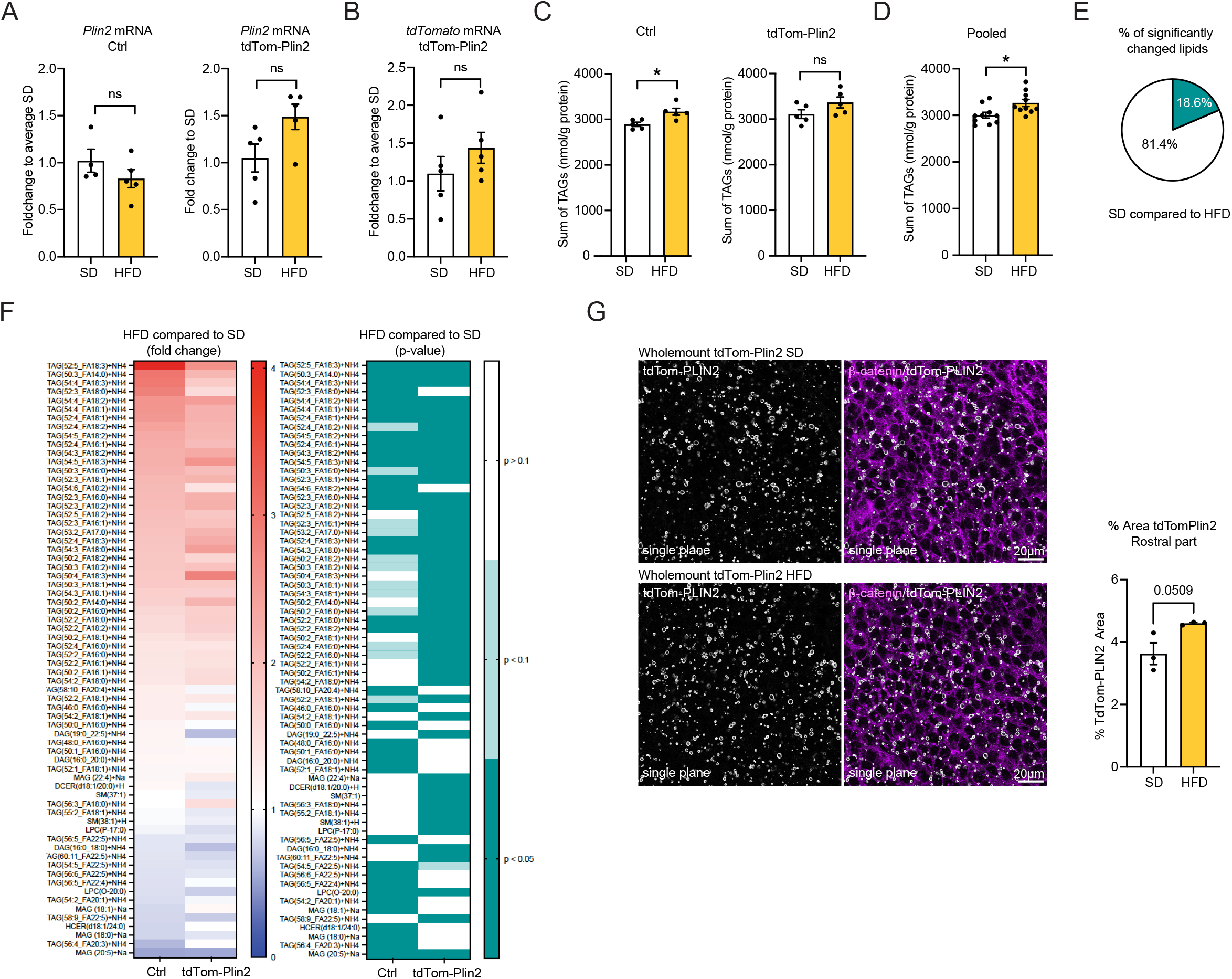
Short-term high fat diet increases the total TAG levels in the brain and leads to slightly increased LDs in the wall of the lateral ventricles. **A)** Analysis of mRNA expression by RT-qPCR show no significant difference in *Plin2* due to the HFD in neither Ctrl mice nor tdTom-Plin2 mice. (n=4-5 mice per group, fold change +/- SEM). **B)** *tdTomato* was also not altered on the gene expression level in SD or HFD tdTom-Plin2 mice (n=4-5 mice per group, fold change +/- SEM). **C** Lipidomic analysis revealed an increase in brain TAGs upon HFD compared to SD, reaching statistical significance for the Ctrl mice (n=4-5 mice/group, mean +/- SEM) and a trend for the tdTom-Plin2 mice. **D)** When both genotypes are pooled, TAGs are significantly increased in HFD fed mice compared to SD mice. (n=9-10 mice per group, mean +/- SEM). **E)** Out of 366 analysed lipids in Ctrl and tdTom-Plin2 brains by lipidomics, 18.6 % were significantly different when comparing SD to HFD. **F)** The majority of the significantly changed brain lipids were TAGs. When looking at their fold changes, both Ctrl and tdTom-Plin2 brains showed a coherent pattern in terms of increased and decreased lipid species as a reaction to the HFD (left heatmap shows fold change to SD, right heatmap shows the corresponding p-values). **G)** Quantification of the area covered by tdTom-Plin2 signal in whole mount preparations of the lateral ventricle wall shows increased LDs with HFD. Representative single planes, ý-catenin outlines the cell membranes of ependymal cells. (n=3 mice per group, mean +/- SEM). Asterisks indicate the following p-values: *< 0.05. ns= non-significant.

We next assessed if the increase in TAGs was evident on the LD level. We used wholemounts of the lateral ventricles of heterozygous tdTom-Plin2 mice on SD or HFD and analysed the area covered by the tdTom-Plin2 signal. Wholemounts provide a clear view of LDs in ependymal cells and NSPCs in both SD and HFD mice (Fig. S5D). Segmentation of the wholemounts into several rostral and medial parts of the lateral wall and quantification of the area covered by tdTom-Plin2 showed that especially the rostral parts had increased LDs with HFD (Fig. 6G, S5C and S5E).

Taken together, we show that the brain has a certain capacity to store TAGs, which increases upon a short term HFD. Importantly, we found that the increase in TAGs was accompanied by an increase in LDs in the lateral ventricle walls, demonstrating that at least ependymal cells are increasing LDs with increased lipid availability. Whether other brain regions also react with increased LDs remains to be determined.

### LDs are present in the developing embryonic brain and react dynamically to exogenous lipids

Little is known about the role of lipid metabolism and LDs during embryonic brain development. As both have been shown to play a role in the regulation of adult NSPCs (Madsen et al., 2021), we investigated if LDs were also present in embryonic brains where NSPCs are actively generating new cells. Querying existing scRNA seq data from the developing brain (http://mousebrain.org) (LaManno et al., 2021), we saw that *Plin2* was expressed in several cell type clusters (Fig. 7A). Embryonic NSPCs, also called radial glial cells (RG) were one of the clusters with more abundant *Plin2* expression (Fig. 7A). Moreover, neuroblasts and neurons also expressed *Plin2.* In line with these scRNA seq data, we found a wide distribution of LDs across the entire thickness of the developing cortex in unstained brain sections from tdTom-Plin2 embryos at E14.5, a stage of highly active neurogenesis (Fig. 7B and C). We confirmed by immunohistochemical staining that the tdTomato signal colocalized well with PLIN2 (Fig. S6A). We next looked in more detail at the ventricular/subventricular zone, which can be labelled with the marker Sox2, and where RGs and intermediate progenitors reside. TdTomato intensity in relation to distance from the ventricle was similar throughout the Sox2-positive layer (Fig. 7D), suggesting a homogenous LD distribution across the ventricular/subventricular zone.

**Figure 7.**
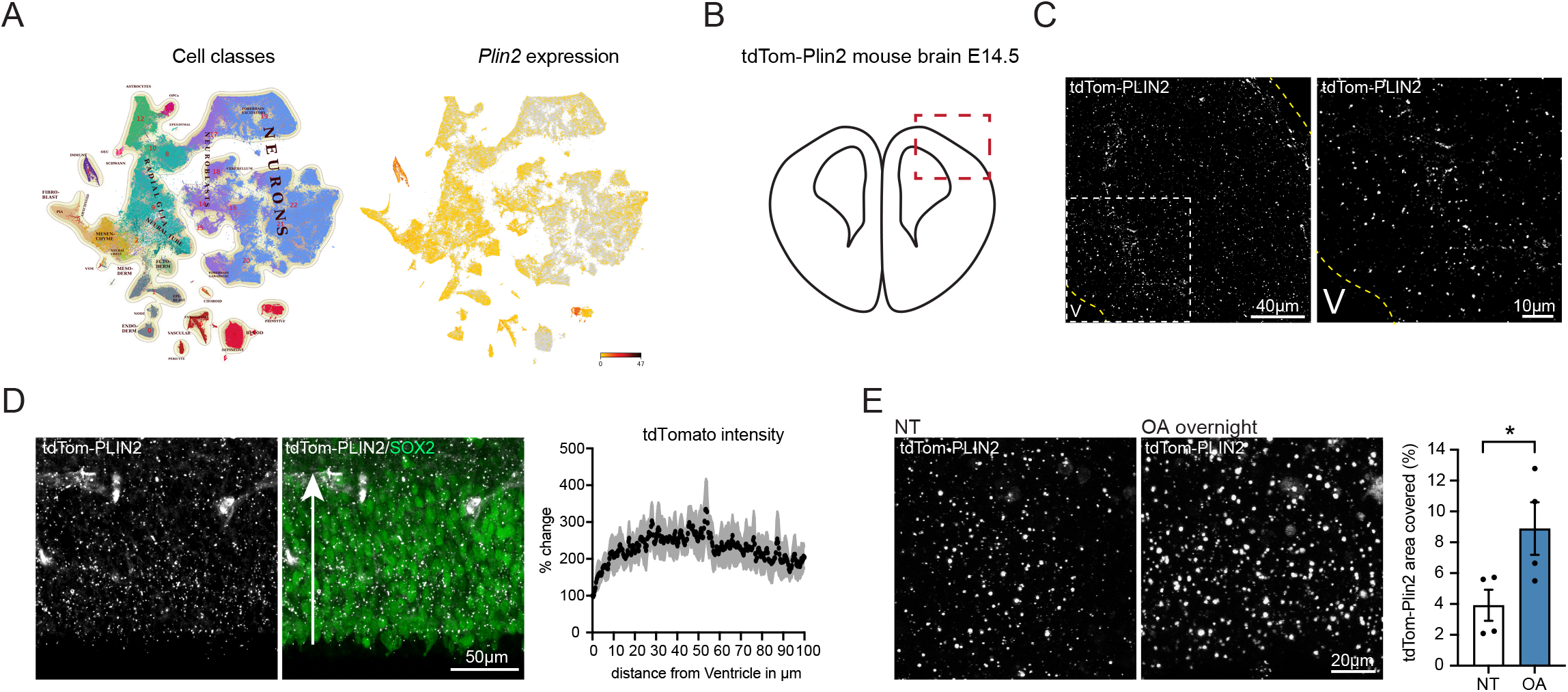
LDs are present in the developing embryonic brain and react dynamically to exogenous lipids. **A)** Querying for *Plin2* expression in scRNA sequencing data from embryonic brain development (http://mousebrain.org) show that *Plin2* is abundant in many clusters, one of them being RG cells. **B)** Illustration of a coronal section from an embryonic brain at E14.5, red square marking the dorsal pallium, where LDs were imaged. **C)** Endogenous tdTom-PLIN2 positive LDs are abundant in the developing cortex at E14.5 in tdTom-Plin2 embryos. (Maximum intensity projection, 25 μm stack, at low and high magnification). **D)** Quantification of tdTomato intensity in relation to distance from the ventricle shows that LDs are homogenously distributed throughout the Sox2-positive layer. Arrow indicates the direction of measurement. **E)** Overnight incubation of OA significantly increases the LD area covered compared to non-treated (NT) sections (n=4 embryos, 2-3 images per embryo, mean +/- SEM).

To investigate if LDs react to exogenous lipids, we incubated acute embryonic brain slices with 0.1 μM oleic acid (OA) overnight. This led to a significant increase in the area covered by LDs (Fig. 7E). To investigate the dynamics of the LDs, we live imaged brain slices from E14.5 tdTom-Plin2 embryos for 2 h in two different conditions: i) sections imaged in artificial cerebrospinal fluid (aCSF), ii) sections imaged in aCSF with 0.1 μM OA after overnight incubation in medium containing 0.1 μM OA (Fig. S6B). Surprisingly, LDs decreased continuously in the non-treated condition during imaging (Fig. S6C), while the decrease in LDs was less prominent after overnight incubation with OA (Fig. S6D). As aCSF contains less nutrients than the medium in which the sections are pre-incubated in, these results suggest that the lipids stored in LDs are used by the cells over time if there are not enough exogenous lipids provided.

In conclusion, we show that LDs are highly abundant in the developing mouse cortex, and that the tdTom-Plin2 mouse also allows visualization of LDs during embryonic brain development. These results also show that LDs react dynamically to exogenous lipids. As a proof-of-concept, live-imaging illustrates the usefulness of the tdTom-Plin2 reporter mouse to study LDs in a dynamic and staining-free manner.

## Discussion

LDs have recently gained new interest as regulated and dynamic organelles, with important functions in health and disease. Here we describe a novel LD reporter mouse, generated by endogenously labelling the LD specific protein PLIN2 with tdTomato using CRISPR/Cas9. We moreover demonstrate the usefulness of this tdTom-Plin2 mouse with various techniques including staining-free detection, scRNA seq and live imaging. We validate the tdTom-Plin2 mouse model under SD and short-term HFD and demonstrate that and increase in LDs are detectable in the livers of the HFD mice. We have further used this mouse model to study LDs in the brain where we show an astonishing abundance of LDs in multiple brain regions and cell types in healthy conditions. We could also show that brain TAGs and LDs increase already after a 4-week HFD. Furthermore, we demonstrate that LDs are present during embryonic brain development and can dynamically react to exogenous lipids, opening new questions on the effect of dietary lipids on brain development and function.

Several tools are available for studying LDs, including classical immunofluorescent staining, lipid dyes such as OilRedO and BD493 and newly developed probes based on lipophilic fluorophores. As they either require fixation prior staining, incubation prior imaging or have limitations in the specificity of LDs (Daemen et al., 2016; Fam et al., 2018; Listenberger and Brown, 2007), better models are needed to study LD dynamics. Existing staining-free approaches comprise specific microscopy techniques, such as Raman microscopy (Yu et al., 2014) or transgenic LD reporters (Targett-Adams et al., 2003), and a few transgenic invertebrate model organisms, such as C.elegans (Liu et al., 2014) and Drosophila (Beller et al., 2010). A very recent addition to the LD imaging toolbox is two LD reporter zebrafish models with fluorescently tagged endogenous Plin2 (Lumaquin et al., 2021) and Plin2/Plin3 (Wilson et al., 2021). However, a mammalian LD reporter model has been missing to this date. Thus, our mouse model adds a valuable tool to the field of LD biology and can be of wide use, as it labels LDs in all cells and tissues expressing Plin2.

Tagging proteins with large fluorophores such as tdTomato always bears the risk of altering the expression or function of the tagged protein. To minimize the impact of the tag, we chose a heterozygous tagging strategy, leaving one *Plin2* allele unaltered. Here we show that endogenous tagging of *Plin2* with tdTomato does not significantly alter the mRNA expression, protein expression or the amount of LDs in NSPCs *in vitro*.

We further show that endogenous tdTom-PLIN2 colocalizes well with stained PLIN2 both in cultured NSPCs and in the brain. Moreover, we report no changes in weight and fat composition of the tdTom-Plin2 mouse under SD and HFD compared to control mice, indicating that the tag does not result in global physiological alterations. Our brain lipidomics analyses further suggests that tdTomato-tagged PLIN2 does not lead to an accumulation or decrease of neutral lipids in the brain, as TAGs, DAGs and MAGs were similar between control and tdTom-Plin2 mice. Furthermore, HFD lead to very similar changes in the brain lipidome in both tdTom-Plin2 and control mice. However, we cannot completely rule out that the tag could exert more subtle changes in LD- or PLIN2 turnover, as the tag could influence access to LDs or protein degradation. This might be to a certain extent tissue- or cell specific and needs to be kept in mind.

Using the tdTom-Plin2 mouse, we show that LDs are found in multiple cell types of the brain under physiological conditions. FACS analyses suggest that approximately 4 % of all the cells in the adult healthy mouse brain have LDs. Crossing the tdTom-Plin2 mouse with neuron- and astrocyte specific GFP reporter lines allowed us to further characterize that around 20 % of all Thy1GFP positive neurons, and almost half of all the Aldh1l1GFP positive astrocytes have LDs. It is important to note that the Thy1GFP mouse sparsely labels neurons throughout the brain (Feng et al., 2000), and Aldh1l1GFP, although being specific to astrocytes, might not be expressed in all astrocytes. As we used whole brain single cell suspensions, it remains to be determined if subtypes of neurons and astrocytes have different LD storage capacities. Moreover, several recent studies suggested LD alterations in the brain with aging or diseased states (Farmer et al., 2020). The tdTom-Plin2 mouse model will therefore provide a powerful tool to study age-related LD accumulation and crossing these mice with neurodegenerative mouse models will enable novel approaches to investigate LD changes under these conditions.

LDs are present in NSPCs *in vitro* (Ramosaj et al., 2021) and in the SVZ *in vivo* (Hamilton et al., 2015). Therefore, we focused on the two main adult neurogenic regions, the SVZ and the DG. In these regions, we show that LDs can be found in neurons, astrocytes, oligodendrocytes, microglia, ependymal cells, endothelial cells and NSPCs. However, not all the cells of a given cell type had LDs. It therefore remains to be investigated if this is due to dynamic intracellular LD fluctuations, or intercellular lipid coupling between cells such as that described between neurons and glia *in vitro* and in *Drosophila* (Ioannou et al., 2019; Liu et al., 2017). Interestingly, almost all NSPCs, as well as astrocytes and neurons *in vitro* have LDs (Ralhan et al., 2021; Ramosaj et al., 2021), whilst only a fraction of these cells has LDs *in vivo.* This might reflect the difference in microenvironment and nutrient availability *in vitro* vs *in vivo;* or it could be due to different stress levels, as LDs can exert protective functions under cellular and oxidative stress (Jarc and Petan, 2019). Thus, these differences highlight the importance of considering the metabolic situation *in vitro* vs *in vivo* when studying the role of LDs. In addition, we found that LD sizes vary a lot among different cell types, with ependymal LDs being much larger than for example LDs in endothelial cells. How these variability in LD size influences the different cell types remains to be determined. As previously suggested for astrocytes and microglia (Kwon et al., 2017; Marschallinger et al., 2020) one reason why not all the cells of a given cell type have LDs *in vivo* could be that the presence of LDs could be linked to distinct subtypes. Indeed, our scRNA sequencing experiment showed a specific LD-containing subpopulation under physiological conditions, based on the presence or absence of tdTom-PLIN2, hence presence or absence of LDs. This population shared some but not all signatures with DAM and LDAM microglia (Chen and Colonna, 2021; Keren-Shaul et al., 2017; Marschallinger et al., 2020), suggesting that DAM and LDAM microglia might already start to develop before disease or aging features appear. It will therefore be interesting to see if this LD-containing population changes with aging and becomes more similar to the previously described disease-associated microglia. As a proof-of-concept, we here show that endogenous fluorescent LDs allow the unbiased separation of cell populations based on LD content, which can then be further explored by downstream analyses such as scRNA sequencing. Our procedure heavily enriched the microglial population, with more than 90% of the high-quality sequencing reads coming from microglia. This is most likely due to their resistance to stresses such as mechanical disruption, followed by FACS and then single cell droplet generation. To fully exploit this novel model, better protocols including whole cell FACS need to be established to cover more cell types.

LDs can accumulate upon altered lipid availability in several tissues, including the liver (Seebacher et al., 2020), and we demonstrate that an increase in LDs can indeed be detected in the liver of tdTom-Plin2 mice after a short term HFD. Surprisingly, we also found a significant increase in TAGs in the brain of HFD fed mice, indicating that even a short exposure to HFD can have an impact on TAGs in the brain. Several studies have addressed brain lipidome alterations upon diet changes, however, as the HFD exposure was much longer in these studies or the diet composition was different (Fitzner et al., 2020; Lee et al., 2018), it is difficult to directly compare those results. Our findings, showing that LDs in the rostral part of the lateral ventricle are increased, suggest that at least certain brain cells can directly react to increased lipid availability with increased lipid storage. This is in line with a study by Ogrodnik and colleagues who looked at how a 2-4-month long exposure to HFD affected the brain of 8-month-old mice. They report an accumulation of PLIN2 in the walls of the lateral ventricles (Ogrodnik et al., 2019), accompanied by a decrease in neurogenesis. Whether and how exactly these two phenomena are linked remains to be further explored.

Relatively little is known about the role of lipid metabolism in brain development. A recent publication showed that *de novo* lipogenesis, which has been found crucial for adult NSPCs (Knobloch et al., 2013), is also critically required in NSPCs during early mouse brain development (Gonzalez-Bohorquez et al., 2022) and human brain development (Bowers et al., 2020). As *de novo* lipogenesis directly influences LD accumulation in adult NSPCs (Ramosaj et al., 2021), we wanted to exploit the tdTom-Plin2 mouse model to address LDs during brain development. We could show that LDs are highly abundant in the developing mouse cortex at E14.5 and can be seen in RGs, in line with scRNA sequencing data of the developing brain (LaManno et al., 2021; Telley et al., 2019). Their high abundance might be linked to high *de novo* lipogenesis, as suggested by the high expression of *Fasn* at this stage of development (http://mousebrain.org, (LaManno et al., 2021). Interestingly, overnight incubation with OA significantly increased LDs, illustrating that external lipids can influence LD abundance. This is in line with a study on cholesterol metabolism during mouse brain development, which showed that LDs increased when cholesterol metabolism was disrupted, potentially due to an increased uptake of lipoproteins from the circulation (Saito et al., 2009). Future studies will have to show if there is a regional, temporal and cell type specific LD abundance and whether manipulation of LD build-up or breakdown can affect brain development. Interestingly, using embryonic mouse brain sections, we demonstrate that the endogenous LD label allows live imaging of LDs in living tissues *ex vivo*. While the tracking of individual cells and the detailed changes of LDs within cells over time will require more work, this illustrates the usefulness of this mouse model for studying the dynamics of LDs.

Overall, we show that the tdTom-Plin2 reporter mouse provides an important novel tool to study LDs and their dynamics. As PLIN2 is ubiquitously expressed, this model is not only interesting for the neuroscience community but can also be used for studying other tissues that contain LDs under physiological or diseased conditions.

## Author contribution

S. M. designed, performed, analysed and interpreted experiments. A. C. D. performed SVZ whole mount experiments and characterized the SVZ. C. C. designed and performed live imaging experiments together with S. M. ScRNA seq experiment were performed and analysed by S. M., E. M., M. K. and L. T.. V.M. performed and analysed metabolic measurements and helped with *in vitro* work. F. B. performed liver experiments and analysed whole mount data. D. J. and F. D. provided resources and interpreted data. M.K. developed the concept, designed, performed, analysed and interpreted experiments. S. M. and M. K. wrote the manuscript, with input from all authors.

## Acknowledgements

We would like to thank A. Denoth Lippuner and Sebastian Jessberger for kindly providing plasmids and input used for the CRISPR approach, R. C. Paolicelli for providing the Thy1GFP mouse and helpful input on the microglia dataset, and P. Bezzi for providing the Aldh1l1GFP mouse. We furthermore thank D. Tavel, M. Niquille and V. Scandella for technical help. We would like to thank the Flow Cytometry facility at University of Lausanne, especially D. Labes for technical help and help with analysis, the Metabolomics Unit at University of Lausanne, specifically T. Taev for technical help and H. Gallart-Ayala for help with analysis. Finally, we thank the Center of PhenoGenomics, EPFL, specifically R. Combe for technical help. This work was supported by funding from the University of Lausanne and the Swiss National Science Foundation (grant # 31003A_175570, M.K), University of Basel and European Research Council Advanced Grant (No 789328, F.D.).

## Material and methods

### Generation of tdTom-Plin2 mice

#### Single guide RNA (sgRNAs) plasmid

sgRNAs targeting the N-terminus of *Plin2* were designed using https://www.addgene.org/crispr/reference/. Oligos (100 μM, Microsynth) were phosphorylated and annealed in a reaction with Oligos (1:10), T4 Polynucleotide Kinase (1:10, #MO201S, New England Biolabs), T4 Ligase reaction buffer (#B0202S, New England Biolabs) with 10 mM 10x Adenosine 5’-Triphosphate (1:10, #P0756S, New England Biolabs). The reaction was incubated in a thermocycler for 37 °C for 30 min, 95 °C for 5 min and then ramped down to 25 °C, 5 °C at a time. The sgRNA plasmid backbone, pSpCas9 (BB)-2A-Puro (PX459) v2.0 (Ran et al., 2013), was digested for 4 h at 37 °C using the restriction enzyme BbsI (#R3539S, New England Biolabs) and run on a 1 % agarose gel for 3.5 h at 100 V. The digested band was cut out and extracted from the gel using QIAquick Gel Extraction Kit (#28704, Qiagen). The sgRNA was then ligated into the backbone by incubation of 10 ng backbone, annealed and phosphorylated sgRNA (1:10) and T4 Ligase reaction buffer (1:10, #B0202S, New England Biolabs) with 10 mM 10x Adenosine 5’-Triphosphate (#P0756S, New England Biolabs) for 30 min in RT.

#### tdTomato plasmid

Mouse *Plin2* homology arms were obtained by extracting DNA from mouse primary mouse NSPCs from the SVZ using DNeasy Blood & Tissue Kit (#69506, Qiagen). The sequences covering the sgRNA target site were extracted from the genomic DNA using PCR, run on a 1 % agarose gel and gel extracted using QIAquick Gel Extraction Kit (#28704, Qiagen). *tdTomato* and *neomycin* were cloned out from H3.1-miCOUNT (Denoth-Lippuner et al., 2020) using PCR, run on a 1 % agarose gel and gel extracted using QIAquick Gel Extraction Kit (#28704, Qiagen). The plasmid backbone, pFA6 (Janke et al., 2004) was digested using the restriction enzymes SacII (#R0157S, New England Biolabs) and HindIII (#R3104T, New England Biolabs). All fragments were cloned together using the Gibson Assembly^®^ Master Mix (#E2611S, New England Biolabs), therefore all primes used for amplifying the fragments were designed to have an overhang to align to the adjacent fragment in the plasmid. The reverse primer for cloning neomycin has a T2A sequence inserted in the overhang linking to *tdTomato*, and the reverse *tdTomato* primer have a GGGGS linker inserted in the overhang linking to *Plin2*. Mutations in the sgRNA target site of the plasmid were introduced and *neomycin* was removed using Q5 Site-Directed Mutagenesis Kit (#E0554S, New England Biolabs).

Once the plasmids were generated, they were sent to PolyGene transgenetics (Switzerland) for the generation of the mouse. In brief, ESCs were CRISPRed and grown as colonies. Selected colonies of ESCs were harvested and DNA was extracted. ESCs were screened for the correct insertion of tdTomato using PCR combined with sequencing of the PCR product. The selected correct ESC clone was then screened for the top three most likely off targets (*Grin2b, Sumf1* and *Gpc6*), based on the online prediction tool: https://cm.jefferson.edu/Off-Spotter/, using PCR and sequencing of the PCR product. The most promising clone of ESCs were then selected for blastocyst injection to generate chimeras. The chimeras were bred and offspring with germline tdTomato insertion was selected.

### Transformation

5-alpha Competent E. coli (High Efficiency) (#C2987I, New England Biolabs) were thawed on ice, 1 μl containing 1 pg-100 ng DNA was added and incubated on ice for 30 min. The bacteria were then heat shocked at 42 °C for 30 s and then placed back on ice for 5 min. 950 μl SOC outgrowth medium (#B9020S, New England Biolabs) was added and sample was incubated on rotation at 37 °C for 1h. 100 μl bacterial suspension was plated on agar plates with ampicillin (100 μg/ml) and incubated at 37°C overnight. Single colonies were picked for further amplification.

### Plasmid amplification

Amplification of plasmids was done using QIAprep Spin Miniprep Kit (#27106, Qiagen) and EndoFree Plasmid Maxi Kit (#12362, Qiagen) according to the manufacturer’s instructions.

### Animals

TdTom-Plin2 mice were generated as described above. C57/Bl6 female mice used for timed mating were purchased from Janvier (France). The Nestin-GPF mouse strain is originally described by Yamaguchi and colleagues (Yamaguchi et al., 2000). The Thy1GFP mice is originally described by Feng and colleagues (Feng et al., 2000) and the Aldh1l1GFP mouse by Gong and colleagues (Gong et al., 2003). All mice were kept under standard conditions in ventilated cages with ad libitum food and water.

Mice fed a HFD (60% energy from fat, SAFE® Complete Care Competence, irradiated #292HF) were housed in the standard conditions, but fed a HFD for 4 weeks starting at the age of 8 weeks. Body weight was monitored on weekly basis, EchoMRI™ was performed at the start and endpoint of the HDF fed mice and the control group fed standard chow diet (#SAFE-150, SAFE® Complete Care Competence). All experiments including animals were carried out in compliance with the Swiss law after approval from the local authorities.

### Collection of mouse tissue

#### Perfusion

Mice were anesthetized by i.p. injection of pentobarbital (150 mg/kg) and intracardially perfused with ice cold 0.9 % saline until no blood remained, then perfused with ∼ 30 ml ice cold PFA, 4 % followed by another ∼10 ml 0.9 % saline. Brain and body were collected and post fixed in ice cold 4 % PFA overnight. Mice exposed to HFD were only perfused with 40 ml 0.9 % ice cold saline, one brain hemisphere and part of the liver was post fixed in 4 % PFA overnight, and the other hemisphere and part of the liver were snap frozen. Serum was collected from mice exposed to HFD by aspiring blood from the ventricle of the heart just before the saline perfusion was started. The blood was transferred to K-EDTA tubes, inverted 5 times and stored on ice for 5-20 min before centrifugation at 1000 x g for 10 min (4 °C). The serum fraction was gently collected with a pipette and transferred to a fresh Eppendorf tube and snap frozen.

#### Subventricular zone whole mount

The subventricular zone whole mount was performed as previously described (Mirzadeh et al., 2010, Doetsch and Alvarez-Buylla, 1996). In brief, mice were sacrificed by decapitation and the lateral walls of the lateral ventricle were dissected out and post fixed in 4 % PFA overnight. The ventricle walls were washed 3 times in 0.1 M PBS before immunohistochemical staining.

#### Sample collection and preparation for lipidomics, RT-qPCR and Western blot

The liver and one snap frozen hemisphere (without olfactory bulb) were homogenized using Evolution & Cryolys® Evolution (#P000062-PEVO0-A.0 & #P000671-CLYS2-A.0, Precellys®), Stainless Metaltube (#P000952-LYSK0-A.0, Precellys®) and 6.8 mm ceramic bead (#P000931-LYSK0-A.0, Precellys®). All sample handling was done in liquid nitrogen. The frozen tissue was placed in a metal tube together with the ceramic bead. The tube was placed in the cryogrinding machine and homogenized at 8800 rpm for 5 sec. The homogenized brain powder was aliquoted for lipidomic analysis, RT-qPCR analysis and western blot. The liver powder was aliquoted for RT-qPCR and western blot.

For the expression analysis of Plin2 in different organs, 3 tdTom-Plin2 het male mice were intracardially perfused with ice cold 0.9 % saline until no blood remained. Organs were dissected and immediately frozen on dry ice. Tissues were mechanically disrupted by passing them through syringes with decreasing needle size (G20 and G25) in RLT lysis buffer supplemented with beta-mercaptoethanol, and RNA extraction was performed with the RNeasy mini kit (#74134, Qiagen) according to the manufacturer’s instructions.

### Brain cell extraction

#### Neural stem/progenitor cell extraction

Adult mouse NSPCs from the SVZ of three 8-week-old female mice were isolated using papain-based MACS Neural Tissue Dissociation Kit (#130-092-628, Milteny) and the GentleMacs Dissociator (Milteny) according to the manufacturer’s instructions.

In brief, mice were shortly anaesthetized with isoflurane, followed by decapitation. SVZ were micro-dissected, and a single cell suspension was generated using the papain-based MACS Neural Tissue Dissociation Kit (#130-092-628, Milteny), according to the manufacturer’s instructions. The single cell suspension was followed by myelin removal using the MACs myelin removal beads (#130-096-731, Milteny,) and a QuadroMACS Separator (#130-090-976, Milteny) according to the manufacturer’s instructions. The obtained cells were cultured as neurospheres in DMEM/F12/GlutaMAX (#31331-028, Gibco) with B27 (#17504044, Gibco), 20 ng/ml human EGF (#AF-100-15, PeproTech), 20 ng/ml human basic FGF-2 (#100-18B, PeproTech), and 1x PSF (#15240062, Gibco). Medium was changed every 2-3 days. The neurospheres were expanded for 5 passages to remove progenitors and other proliferating cells. After 5 passages, cells were changed to the following culture medium: DMEM/F12/GlutaMAX (#31331-028, Gibco), N2 (#11520536, Gibco), 20 ng/ml human EGF (#AF-100-15, PeproTech), 20 ng/ml human basic FGF-2 (#100-18B, PeproTech), 5mg/ml Heparin (#H3149-50KU, Sigma) and 1x PSF (#15240062, Gibco). All the *in vitro* experiments were done on pooled NSPCs from 3 mice, passage 7-15.

#### Whole brain cell extraction

Protocol adapted from (Mattei et al., 2020). In brief, mice were anesthetized by i.p. injection of pentobarbital (150 mg/kg) and intracardially perfused with 30 ml cold PBS (pH 7.3-7.4). The brain was collected and finely chopped with a few drops of Hibernate A (Hibernate®A minus phenol red, BrainBits) before complete dissociation in 1.5 ml Hibernate A in a 1 ml douncer (loose pestle). The cell suspension was filtered twice in a prewet 70 μm cell strainer (#15370801, Fisher Scientific), the rubber tip of a 1ml syringe was used to facilitate the filtering. The cell suspension was transferred to a precooled 15 ml tube and centrifuged at 500 x g for 6 min. The supernatant was aspired, and the cell pellet was resuspended in 2 ml cold PBS (pH 7.3-7.4), before adding 1 ml isotonic percoll in PBS (pH 7.3-7.4) (#GE17-0891-01, Sigma-Aldrich). The suspension was mixed well before 4 ml cold PBS (pH 7.3-7.4) was gently added on top of the cell suspension. The layered sample was centrifuged at 3000 x g, 10 min. The supernatant and the myelin disk were gently aspired. The cell pellet was gently washed by adding 4 ml cold PBS (pH 7.3-7.4) and gently tilting the tube back and forth before centrifuging at 450 rcf, 10 min. Cell pellets used for FACS were dissolved in 500 ul EDTA-DPBS (#E8008, Merck Millipore), cell pellets used for scRNA sequencing were dissolved in 500 μl PBS (pH 7.3-7.4).

#### ScRNA seq and analysis

After sorting, cells were immediately processed according to the 10X Chromium protocol. Briefly, an appropriate volume of each cell suspension containing cells from tdTomato positive and tdTomato negative conditions were combined with 10X Chromium reagent mix and samples were loaded into two separate lanes. Cell capture, lysis, mRNA reverse transcription, cDNA amplification and libraries were performed following 10X Genomics Chromium dual indexing Single Cell 3’ V3.1 reagent kit instructions. Libraries were then multiplexed and sequenced according manufacture recommendations with paired-end reads using a HiSeq4000 platform (Illumina) with an expected depth of 100’000 reads per single cells. All the sequencing experiments were performed within the Genomics Core Facility of the University of Lausanne. Alignment of sequenced reads to the mouse genome (GRCm38) and filtered gene– barcode matrices were realized by running Cell Ranger Single-Cell Software Suite v5.0.1 (10X Genomics). The cell ranger count function was used to generate filtered gene/cell expression UMI corrected matrices by selecting probable cells and removing empty lipid droplets. To filter only high-quality cells, we applied selection based on mitochondrial genes percentage (<10 %) and number of genes per cell (>500 genes), doublets cells were removed using Scrublet package. After applying these filters, 845 cells for “tdTomato positive” condition and 784 cells for “tdTomato negative” condition were kept for further analysis. The cells were then annotated using reference single cell atlas, and microglial cells were kept for following analysis (812 microglia for “tdTomato positive” condition and 766 microglia for “tdTomato negative” condition). The raw count matrix was normalized and scale using SCTransform procedure from Seurat package. For UMAP visualization, dimensionality reduction was performed using standard function from Seurat. We first adopted a graph-based clustering approach using “FindClusters” function with a resolution of 0.3. Differentially expressed genes were identified based on their weight, resulting from differential pairwise expression analysis using Seurat “FindAllMarkers” function with default parameters. The identified gene candidates for each condition were interrogated for statistically significant gene ontologies analysis using the free online tool Metascape (metascape.org).

#### Cell culture

NSPCs were grown on uncoated plastic cell culture dishes for expansion (#430167, Corning, 100 mm * 20 mm, TC-treated). Cells used for experiments were plated on glass coverslips (#10337423, Fisher) or standard plastic cell culture dishes coated with poly-L-ornithine (#P3655, Sigma) and laminin (#L2020-1MG, Sigma). Proliferating NSPCs were kept in DMEM/F12/GlutaMAX (#31331-028, Gibco) complemented with N2 (#11520536, Gibco), 20 ng/ml human EGF (#AF-100-15, PeproTech), 20 ng/ml human basic FGF-2 (#100-18B, PeproTech), 5 mg/ml Heparin (#H3149-50KU, Sigma) and 1x PSF (#15240062, Gibco). Medium was changed every 2-3 days. EdU pulse was done by incubating the cells with EdU (1:1000) (2.52 mg/ml, Click-iT Plus EdU Alexa Fluor 647 Imaging Kit, # 15224959, Invitrogen) for 1 h at 37 °C before fixation with 4 % PFA.

#### Metabolic measurements

Oxygen consumption rate (OCR) and extracellular acidification rate (ECAR) of NSPCs was measured using the Seahorse XF Cell Energy Phenotype Test Kit (Agilent, # 103325-100), according to the manufacturer’s instructions. Measurements were performed on a Seahorse XFe96 analyzer (Agilent) in a 96-well plate format. In brief, 40′000 cells were plated on coated Seahorse 96-well plates and left to attach overnight, in 80 μl proliferation medium per well. Cells were then washed 2 times with XF DMEM assay medium (Agilent, #103575-100), supplemented with glucose (Agilent, 103577-100 Seahorse XF 1.0 M glucose solution), glutamine (Agilent, 103579-100 Seahorse XF 200 mM glutamine solution) and pyruvate (Agilent, 103578-100 Seahorse XF 100 mM pyruvate solution) with final concentrations as following: glucose 10 mM, glutamine 2 mM, pyruvat 1 mM. Before starting the measurements, NSPCs where incubated for 1 h in a non-C02 but humidified 37 °C incubator (Agilent) in the same XF assay medium as described above, including 20 ng/ml human EGF and 20 ng/ml human basic FGF-2. Baseline OCR and ECAR were measured over 3 timepoints (15 min). Oligomycine (1 µM) and FCCP (1 µM) were injected into each well using the Seahorse Cartridge system and OCR and ECAR measured immediately after.

At the end of the measurements, live Hoechst 33342 was injected in each well and cells were incubated for 15 min to stain the nuclei. Cells were fixed with 2% PFA and images of the entire wells were taken with a Thunder microscope, 5x objective. Area covered by the Hoechst signal was calculated using ImageJ and this value was used for normalization of the OCR and ECAR values of each well. For the baseline, the 3rd measure (after 14.5 min) was used. For the metabolic capacity, the difference between the first measure after the oligomycin and FCCP injection and the baseline was calculated. 12-15 replicates per condition were averaged and the experiments repeated 3 times.

#### Lipidomic analysis

Lipidomic analysis of brains (analysing around 1200 lipid species belonging to five different subclasses including TAGs, DAGs, MAGs, CEs, sphingolipids, glycerophospholipids, free fatty acids) was done as previously described (Medina et al., 2020). In brief, lipids were extracted from 25 mg brain tissue sample prepared as described above. The extracted lipids were then analysed using Hydrophilic Interaction Liquid Chromatography coupled to tandem mass spectrometry (HILIC - MS/MS) in positive ionization mode using an TSQ Altis triple quadrupole instrument (Thermo Fisher Scientific) followed by chromatographic separation carried out on an Acquity BEH Amide, 1.7 μm, 100 mm × 2.1 mm I.D. column (Waters). Raw LC-MS/MS data was processed using the MultiQuant Software (version 3.0.3, Sciex technologies). Relative quantification of metabolites was based on Extracted Ion Chromatogram areas for the monitored MRM transitions. The obtained tables (peak areas of detected metabolites across all samples) were exported to “R” software (http://cran.r-project.org/) where the signal intensity drift correction was done within the LOWESS/Spline normalization program followed by noise filtering (CV (QC features) > 30 %) and visual inspection of linear response. Lipid concentrations were normalized to sample protein concentration.

#### FACS

Isolation of NSPC from the SVZ and hippocampi from 3 male NesGFP x tdTom-Plin2 mice, one NesGFP mouse and one tdTom-Plin2 mouse was performed as described above. But instead of plating cells for propagation, they were dissolved in EDTA-DPBS (#E8008, Merck Millipore), and stained with the viability markers DAPI (1 μg/ml) and RedDOT (1:500, #40060-T, Biotium) before being sorted on a FACSAria II (BD Biosciences).

Whole brain cell isolation from 3 male Aldh1l1GFP x tdTom-Plin2 mice, 3 male Thy1GFP x tdTom-Plin2 mice, one male Aldh1l1GFP mouse, one male Thy1GFP mouse and 2 male tdTom-Plin2 mice were sacrificed by decapitation and performed as described above but without anesthetization and perfusion. Cell pellets were dissolved in EDTA-DPBS and stained with DAPI and RedDOT before being sorted on a FACSAria II.

### Immunocytochemical and immunohistochemical staining

Cells were fixed with 4 % PFA (37 °C) for 15-30 min at RT, washed 2 times with PBS and stored at 4 °C. Antibodies and their dilution factor are as following: Actin mouse (1:5000, Sigma-Aldrich A2228), beta-catenin rabbit (1:200, Cell signaling #9587S), beta-catenin mouse (1:200 BD 610154), CD31 rat (1:100 BD 550274), DCX guinea pig (1:500 Merck Millipore AB253), EGFR goat (1:100 R&D BAF1280), EGFR rabbit (1:100 Abcam ab52894), GFAP chicken (1:600 Millipore AB5541), GS rabbit (1:100 Abcam ab73593), IBA1 rabbit (1:1000, Wako Chemicals 019-19741), NG2 rabbit (1:100 Millipore ab5320), OLIG2 rabbit (1:100, Millipore AB9610), PLIN2 rabbit (1:1000 Abcam ab52356), S100b mouse (1:100, Sigma-Aldrich S2532), SOX2 goat (1:500, R&D Systems AF2018-SP), tdTomato goat (1:500, Sicgen AB8181-200).

Immunocytochemical staining were performed as described (Listenberger and Brown, 2007). In brief for saponin protocol, cells were incubated in blocking buffer (1.5 % Glycine, 3 % BSA, 0.01 % Saponin in PBS) for 45 min in RT. Followed by incubation in primary antibody diluted in antibody diluent (0.1 % BSA, 0.01 % Saponin in PBS) overnight, 4 °C. After 3 washes in PBS, cells were incubated in secondary antibody diluted in antibody diluent for at least 1 h in RT, protected from light. Cells were washed 2 times with PBS. Thereafter incubated with DAPI (D9542, Sigma) diluted in TBS for 10 min, followed by one wash in TBS. Coverslips were mounted with a homemade PVA-DABCO-based mounting medium. EdU staining was done using the Click-iT Plus EdU Alexa Fluor 647 Imaging Kit (#15224959, Invitrogen) according to the manufacturer’s instructions.

Post fixed brains were, if cut on a microtome, incubated in 30 % sucrose overnight before being cut into 40 μm sagittal sections on a microtome. Coronal sectioning was done on a vibratome, generating 25 μm sections. Mice for HFD experiment were embedded in 4 % agar before coronal sectioning. Brains from E14.5 embryos were incubated in 30 % sucrose and cut in 60 μm coronal sections on a microtome. The sections were then stored in cryopreservation solution (25 % ethylene glycol, 25 % glycerol in 0.05 M phosphate buffer) at 4 °C.

Immunohistochemical staining was done as follows. Brain sections were washed two times 5 min in PBS on a shaker before incubation in blocking buffer (0.05-0.25 % Triton X-100, 3 % donkey serum in TBS) for 1 h on a shaker, RT. Sections were then incubated in primary antibody in blocking buffer at 4 °C overnight on a shaker. The brain sections were washed 3 times 10 min with PBS and then incubated in secondary antibody in blocking buffer for at least 1 h on shaker in RT. The sections were then washed one time 10 min in PBS and one time 10 min in TBS before incubation with DAPI in TBS for 10 min. Brain sections were then mounted on Superfrost (#10149870, Fisher Scientific) and homemade PVA-DABCO-based mounting medium.

When detecting LDs, staining was performed according to the previously mentioned procedure, but blocking was done using the saponin blocking buffer (1.5 % Glycine, 3 % BSA, 0.01 % Saponin in PBS) and antibodies were incubated in antibody diluent (0.1 % BSA, 0.01 % Saponin in PBS).

Livers were incubated in 30 % sucrose overnight before being cut into 40 μm sections on a microtome. Staining was performed on free-floating sections using the saponin protocol. BD493 (#11540326, Fisher Scientific, in DMSO) was added 1:1000 in antibody diluent and sections were incubated for 1 h at RT on an orbital shaker. Sections were washed 3 times 10 min in PBS, Nuclei were counterstained with DAPI in TBS, followed by additional 2 times 10 min wash in TBS and sections were mounted as described above. To assess the endogenous tdTomato signal, sections were incubated for 10 min in the blocking buffer from the saponin protocol, nuclei were stained with DAPI followed by 2 times 10 min wash in TBS and sections were mounted as described above.

#### RT-qPCR

NSPC pellets from cultured cells were snap frozen on dry ice. RNA was extracted using RNeasy^®^ plus mini kit (#74134, Qiagen) according to the manufacturer’s instructions. cDNA preparation was done using the SuperScript™ IV First-Strand Synthesis System with oligo-dT primers (#15327696, Invitrogen). RT-qPCR was performed using TaqMan™ Fast Advanced Master Mix (#4444557, Thermo Fischer Scientific) and Applied Biosystems TaqMan Assays. The following probes were used: Plin2 (Mm00475794_m1), tdTomato (Mr07319439), 18S (4333760T), b-Actin (Mm01205647_g), Atgl (Mm00503040), Hsl (Mm00495359_m1), Mgl Mm00449274_m1. Results were analysed using the ddCT method, normalizing the samples to either eukaryotic 18S rRNA or mouse *beta-actin*. Statistical analyses were performed on the dCT values.

#### Western Blot

NSPCs were washed once with PBS, collected, and spun down at 300 x g for 3 min. The supernatant was aspired and the cell pellet was resuspended in RIPA buffer (#R0278-50ML, Sigma-Aldrich) with protease inhibitors (#11873580001, Sigma) followed by incubation at 4 °C for 1 hour. The cell lysate was then centrifuged at 12000 x g for 15 min at 4 °C. The supernatant was collected, and the protein concentration was measured using Pierce BCA Protein Assay Kit (#23227, Fisher Scientific) according to the manufacturer’s instruction. The protein samples were mixed with Laemmli sample buffer (#1610747, Bio-rad) supplemented with beta-mercaptoethanol and denaturated at 95 °C for 5 min before loading on a 12 % PAA precast gel (#4561044, Bio-Rad). The gel was run at 70 V for 15 min and then 100 V for 1 h. The proteins were then transferred to a Amersham^™^ Protran^®^ Western-Blotting-Membrane (#GE10600044, Sigma-Aldrich) at 120 V for 1 h. The membrane was then incubated in blocking solution (5 % milk powder, 0.05 %Triton in TBS) followed by overnight incubation of primary antibody in blocking solution. The membrane was washed 3 times, 10 min, in TBS before incubation of secondary antibody in blocking buffer. The membrane was washed 3 times for 10 min in TBS before being revealed using ECL technology with WesternBright ECL HRP Substrate (K-12045-D20, Witec).

#### Serum analysis

Serum analysis of ALAT and ASAT, Glucose, Total Cholesterol, High Density Lipoprotein, Low Density Lipoprotein, Triglycerides and Free Fatty Acids was one with the Dimension Xpand Plus automated chemistry system (Siemens Healthcare Diagnostics AG) at Center of PhenoGenomics, EPFL (Swizerland).

#### Live tissue preparation and live imaging of LDs in E14.5 brain sections

Time mated TdTom-Plin2 females were sacrificed at day 14.5 after mating through anaesthetisation with isoflurane followed by decapitation. The uterine horns were exposed by caesarean cut and embryo heads were collected in ice-cold Ca^2+^/Mg^2+^-free Hank’s balanced salt solution (HBSS, Gibco). Brains were rapidly dissected out and embedded in 3 % low-melting point agarose (LMP-agarose, #6351.2, Roth). Coronal 250 µm-thick slices at the level of the somatosensory cortex were then cut in ice-cold HBSS on a Vibratome (#VT1000S, Leica) and kept in their surrounding agarose. Slices were then placed in an incubator (37 °C, 5 % CO_2_) for at least one hour of recovery on floating nucleopore track-etched polycarbonate membranes (013 mm, 1 µm pore size, #WHA110410, Whatman) in supplemented DMEM-F12 medium (2 % B27, 2 mM GlutaMAX, 1 % penicillin-streptomycin and 1 % N2). Prior to imaging, slices were labelled with SPY650-DNA (1:1000, #SC501, Spirochrome) for 1 h and CellTrace^TM^ CFSE (1:1000, #C34554, ThermoFisher) for 30 min. Slices were then washed, transferred, anchored and superfused with warm (37 °C) and bubbled (medical oxycarbon) aCSF (125 mM NaCl, 26 mM NaHCO_3_, 10 mM D-Glucose, 3 mM KCl, 1.5 mM MgCl_2_, 1.25 mM NaH_2_PO_4_, 1.6 mM CaCl_2_) in a standard cover glass-mounted chamber (#64-0265, Warner Instruments) on an inverted confocal microscope (Nikon A1r), equipped with oil-immersion 60x objective (1.4 Plan Apo VC H, Nikon). 30 µm-thick stacks (2 µm-stepped) were acquired with a 2x numerical zoom every 5 min with resonant laser scanning in the VZ of the dorsal pallium. For OA experiments, slices were incubated in 0.1 μM OA.

#### Imaging and image analysis

All images used for LD quantification were acquired with a confocal microscope (Zeiss, 780, 800 and 900) or a spinning disk confocal (Nikon Ti2, Yokogawa CSU-W1) with a 40x or 63x objective with digital zoom. Images for assessing cell proliferation *in vitro* was acquired using an epifluorescent microscope (Nikon, 90i) with a 20x objective.

*In vitro* colocalization of tdTomato and PLIN2 was assessed using Fiji (Version 2.0.0-rc-69/1.52p). In brief, the images were pre-processed using “Z-project”, “8-bit conversion”, “Subtract Background”, “Brightness/Contrast”, “Set threshold” and converted to mask. Both tdTomato and PLIN2 was then quantified using “Analyse Particles”, and a mask was created. Colocalization of the masks of the two channels were assessed using “colocalization”. The number of EdU+ cells were also assessed using Fiji. The EdU image was processed as follows “8-bit conversion”, “smooth”, “enhance contrast”, “set threshold”, conversation to mask, “fill holes” and “watershed”. The mask was then quantified using “Analyse Particles”. PLIN2 *in vitro* and *in vivo* was quantified by “Analyse Particles”, after the same pre-processing as for assessing colocalization of tdTomato and PLIN2. The number of cells in all conditions was quantified through manual counting of DAPI positive nuclei.

Area covered of NesGFP and DCX positive cells in the granule cell layer of the DG were quantified using Fiji (Version 2.0.0-rc-69/1.52p). The images were re-processed by “Z-projection”, “8-bit conversion”, “Set threshold” and conversion to mask before area covered were measured and normalised to the DG area defined using DAPI or GFP.

Area covered of tdTom-PLIN2 in wholemount preparations of the lateral ventricles were quantified using Fiji (Version 2.0.0-rc-69/1.52p). In brief, the images were processed using “Color Balance” to remove the beta-Catenin signal, “8-bit conversion”, “Brightness/Contrast”, “Set threshold” and converted to mask. The mask was then divided into 8 zones in which the tdTom-Plin2 signal was quantified using “Measure”. Area covered of BD493 in liver sections was quantified using Fiji (Version 2.0.0-rc-69/1.52p). In brief, the images were processed using “8-bit conversion”, “Subtract Background”, “Set threshold” and converted to mask. BD493 was then quantified using “Analyse Particles” and normalized to the DAPI area. The area covered of DAPI was processed using “8-bit conversion”, “Gaussian Blur”, “Set threshold” and converted to mask before being quantified using “Analyse Particles”. Area covered of tdTom-PLIN2 in liver sections was quantified using Fiji (Version 2.0.0-rc-69/1.52p). In brief, the images were processed using “Brightness/Contrast”, “Subtract Background” and then the mean intensity was quantified using “Measure” and normalized to the DAPI area. All quantifications were done blinded.

#### Statistics

Statistical analyses were performed with Prism (GraphPad) as following. A student t-test was used for comparing two groups. Serum analyses was done using a 2-way ANOVA. Significance was considered for p-values < 0.05. For fold change (FC) analyses compared to a control group, FC values were log2 transformed and a one-sample t-test was performed. The nature of the sampling (“n”) is described in each figure legends. A minimum of n=3 is used for each statistical comparison.

---

**Supplementary Figure 1.**
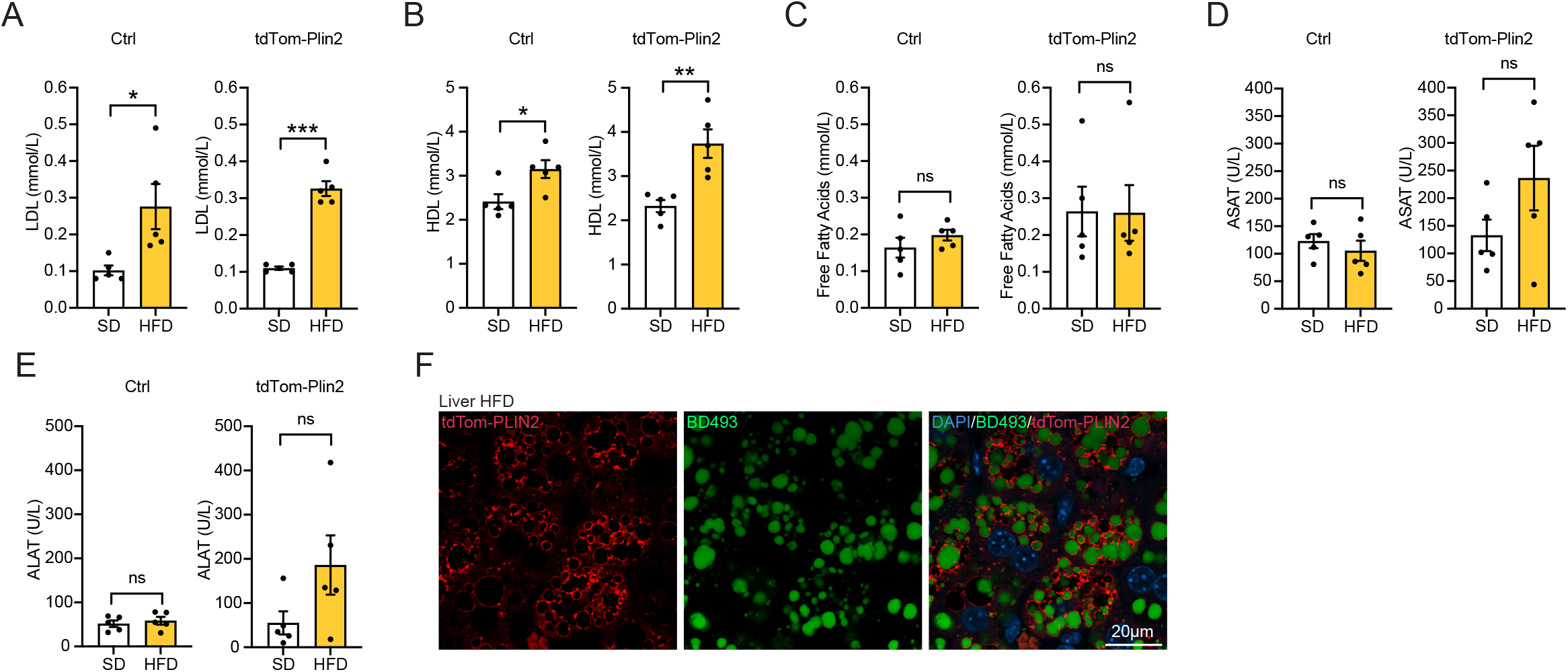
TdTom-Plin2 mice increase fat mass and LDs in the liver upon a short-term high fat diet, related to Figure 2. **A** and **B)** Serum analysis of Ctrl and tdTom-Plin2 mice on SD or HFD show a significant increase in low-density lipoprotein (LDL) cholesterol and high-density lipoprotein (HDL) cholesterol after HFD. (n=5 mice per group, mean +/- SEM). **C)** Free fatty acids in the serum were not significantly changed. **D** and **E**) Blood levels of alanine aminotransferase (ALAT) and aspartate aminotransferase (ASAT), which are used as indicators of liver disease, were not significantly changed with HFD, even though tdTom-Plin2 mice on HFD had a slight elevation in both. **F)** Representative single plane image of a HFD liver of a tdTom-Plin2 mouse, show BODIPY 493/503 (BD493) positive LDs surrounded by tdTom-PLIN2.

**Supplementary Figure 2.**
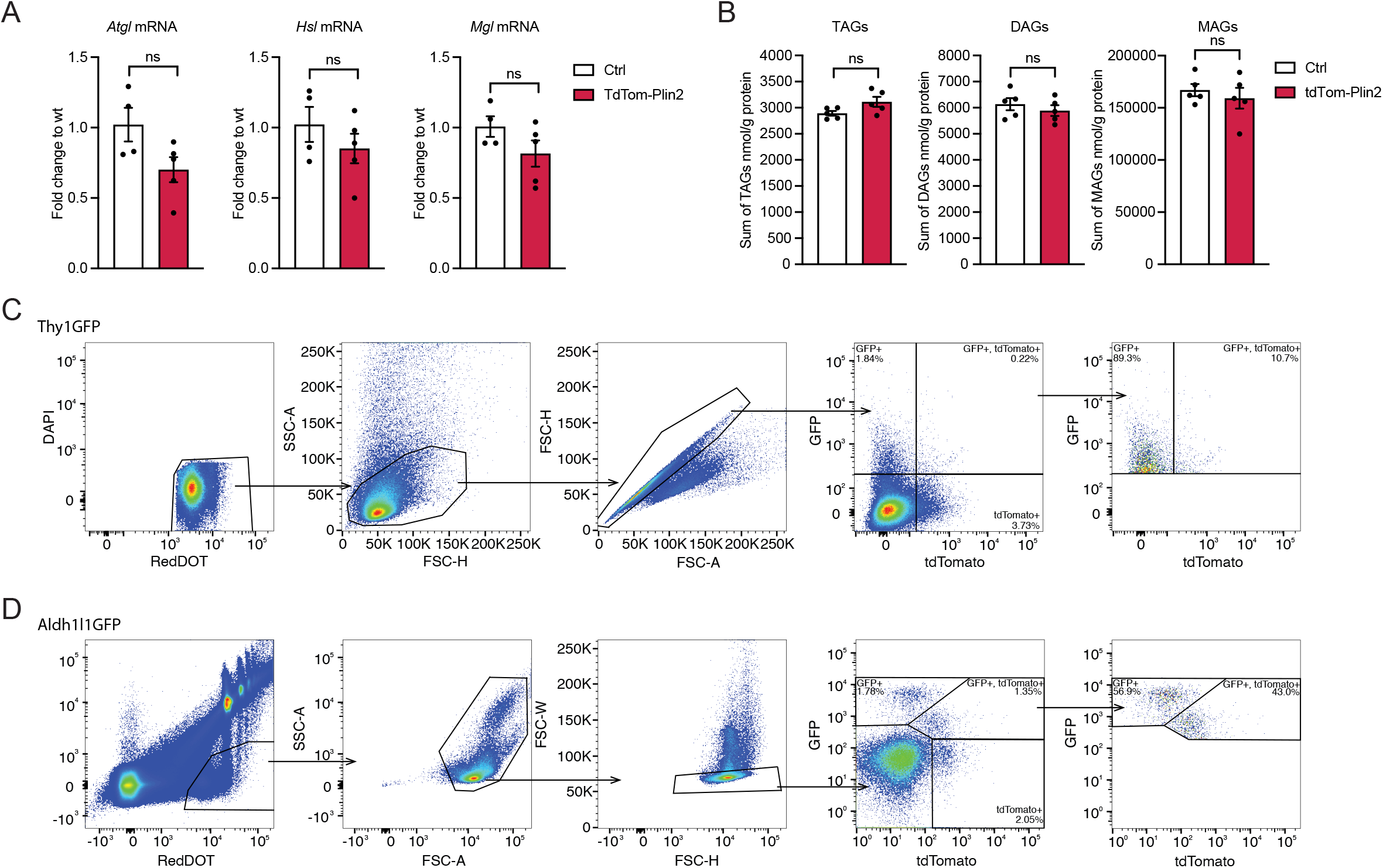
LDs are present in multiple cell types in the healthy adult mouse brain, related to Figure 3. **A)** Analysis of mRNA expression by RT-qPCR show no significant difference in the lipases *Atgl, Hsl* and Mgl in the brain of Ctrl and tdTom-Plin2 mice. (n=4-5 mice per group, fold change +/- SEM). **B)** There are no significant differences in brain TAGs, DAGs and MAGs, measured by lipidomics analysis, between Ctrl mice and tdTom-Plin2 mice. (n=5 mice per group, mean +/- SEM). **C** and **D)** FACS gating strategy for Aldh1l1GFP positive astrocytes and Thy1GFP positive neurons. All samples were first selected on viability based on DAPI and RedDOT staining, followed by sorting based on size and granularity, exclusion of doublets to have a population of viable single cells. These were then sorted based on GFP (cell marker) and tdTomato (tdTom-Plin2) to quantify how many cells have LDs in the different cell populations of the brain.

**Supplementary Figure 3.**
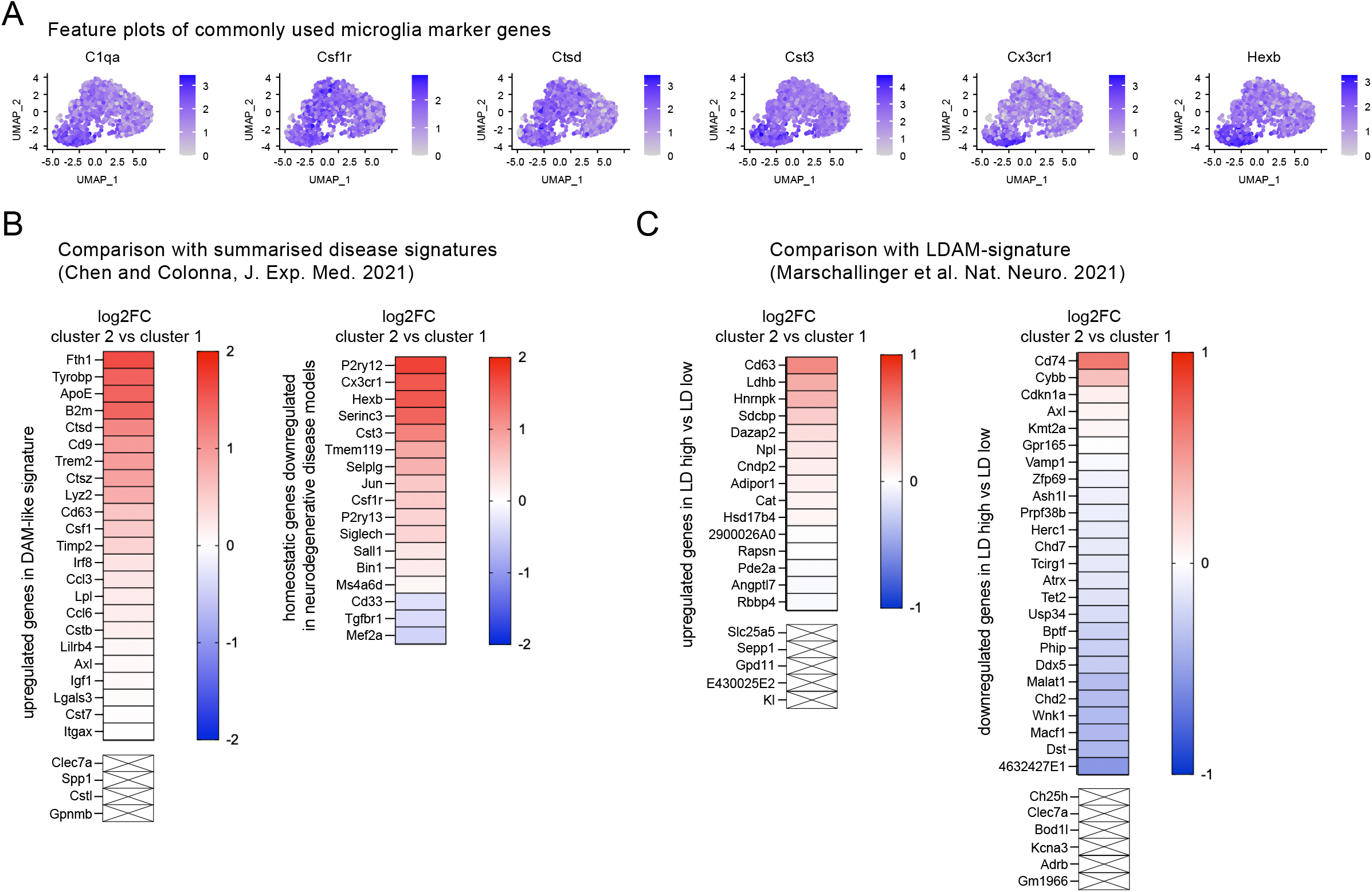
ScRNA sequencing of tdTomato positive microglia reveals a subpopulation with a specific signature under physiological conditions, related to Figure 4. **A)** Feature plots of commonly used microglia marker genes shows the microglial nature of the cells analysed. **B)** Comparison of the gene signature of the tdTom-Plin2 enriched cluster 2 with the summarised disease signatures published by Chen and Colonna (Chen and Colonna, 2021). **C)** Comparison of the gene signature of the tdTom-Plin2 enriched cluster 2 with the LDAM signature published by Marschallinger and colleagues (Marschallinger et al., 2020).

**Supplementary Figure 4.**
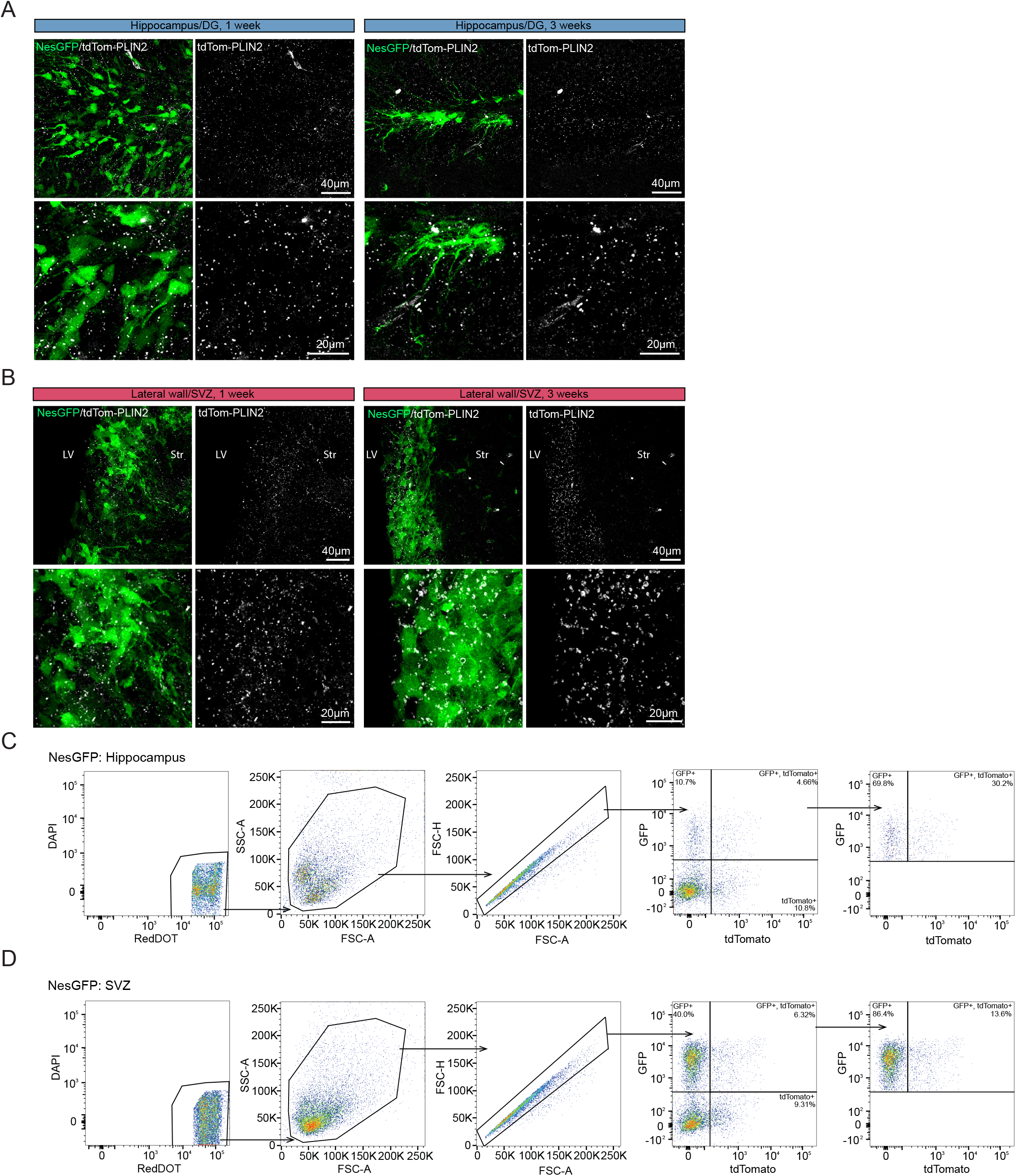
LDs are present in NSPCs and their progeny in the postnatal and adult mouse brain, related to Figure 5. **A** and **B)** Endogenous tdTom-PLIN2 reveals that LDs are abundant the DG and SVZ of both 1 week and 3 weeks old mice. Representative images of non-stained sections showing NesGFP positive NSPCs and tdTom-PLIN2 positive LDs in the DG and SVZ. (maximum intensity projections, 20 μm stacks). **C** and **D)** FACS gating strategy for NesGFP positive cells in the hippocampus and SVZ. All samples were first selected on viability based in DAPI and RedDOT staining, followed by sorting based on size and granularity, exclusion of doublets to have a population of viable single cells. These were then sorted based on GFP (cell marker) and tdTomato (tdTom-Plin2) to quantify how many cells have LDs.

**Supplementary Figure 5.**
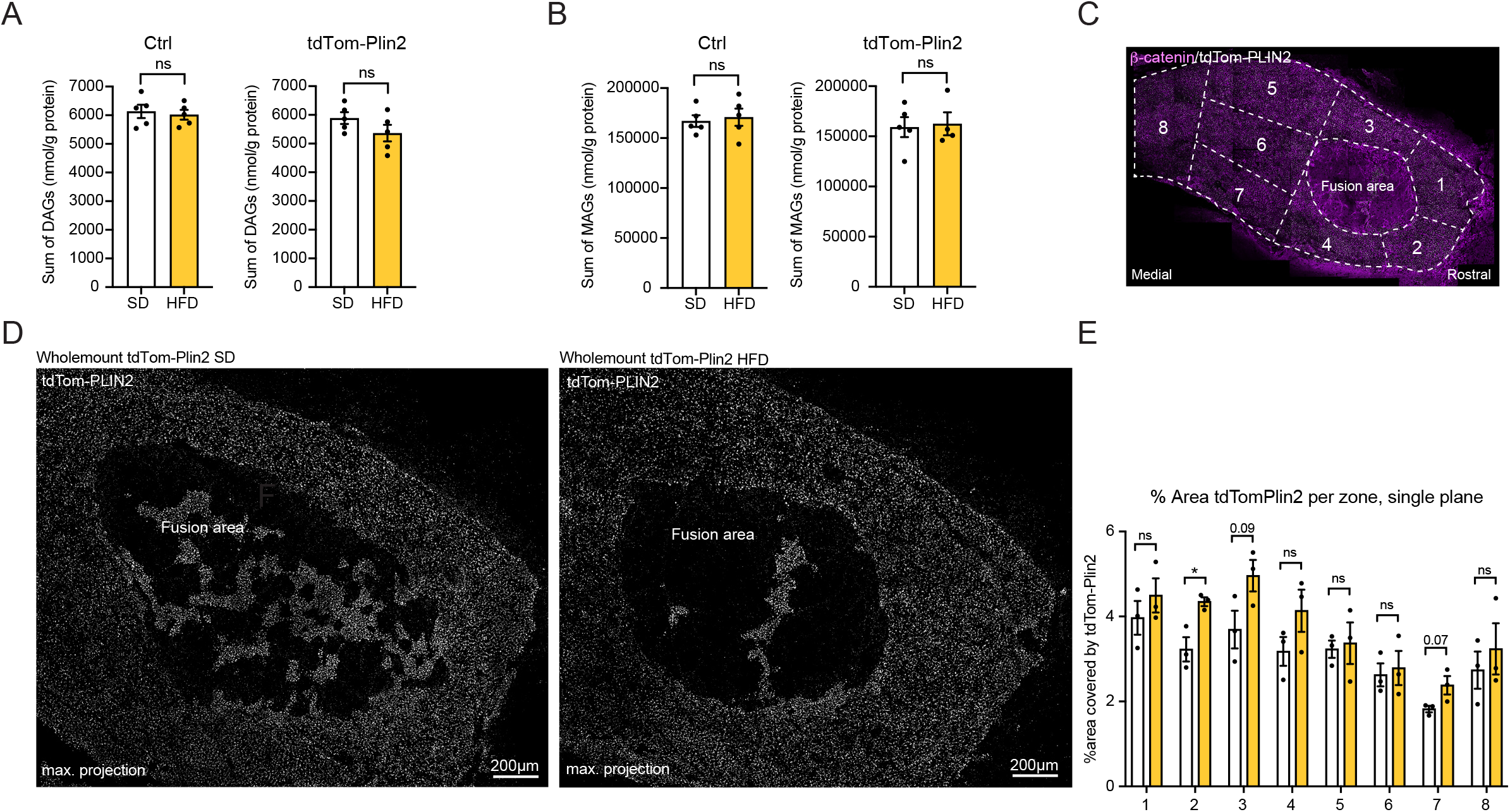
Short-term high fat diet increases the total TAG levels in the brain and leads to slightly increased LDs in the wall of the lateral ventricles, related to Figure 6. **A** and **B**) Lipidomic analysis show no significant difference in DAGs and MAGs after HFD in neither Ctrl or tdTom-Plin2 mice. (n=5 mice per group, mean +/- SEM). **C)** Outline of the zones analysed in the wholemount preparations of SD and HFD tdTom-Plin2 mice. **D)** Wholemount preparation of the lateral ventricle showing LDs stored in ependymal cells and NSPCs in both SD (left panel) and HFD (right panel) mice. **E)** Quantification of the area covered by tdTom-Plin2 in the different zones. (n=3 mice per group, mean +/- SEM). Asterisks indicate the following p-values: * < 0.05; ns= non-significant.

**Supplementary Figure 6.**
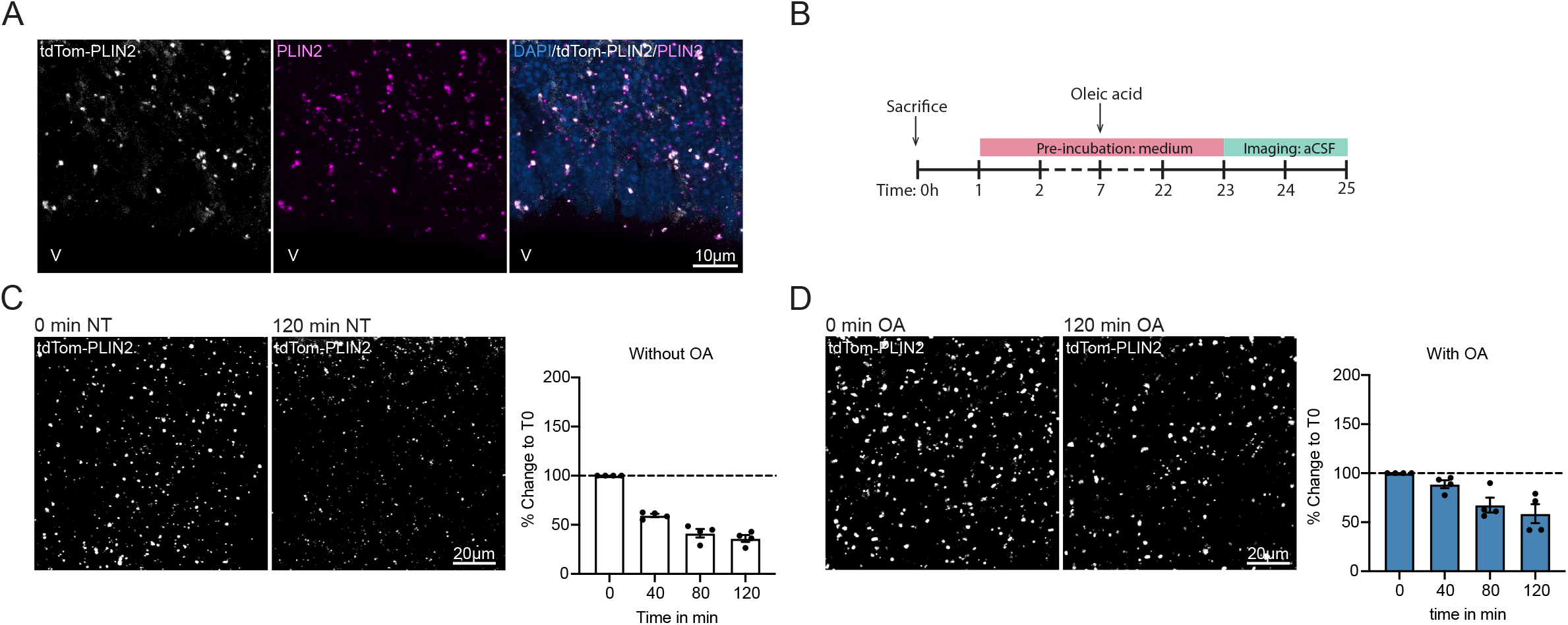
LDs are present in the developing embryonic brain and react dynamically to exogenous lipids, related to Figure 7. **A)** Immunohistochemical staining with PLIN2 shows good colocalization of PLIN2 to tdTom-Plin2. **B)** Timeline of OA treatment before live imaging. **C** and **D)** Representative images showing tdTom-Plin2-positive LDs in NT and OA treated sections at the start point (0 min) and endpoint (120 min) of the live imaging. Maximum intensity projections, 30 μm stacks. (n=4 embryos, 2-3 images/embryo, mean +/- SEM).

## Notes

### Competing Interest Statement

The authors have declared no competing interest.

